# A nanobody-based toolset to monitor and modify the mitochondrial GTPase Miro1

**DOI:** 10.1101/2021.12.10.472061

**Authors:** Funmilayo O. Fagbadebo, Philipp D. Kaiser, Katharina Zittlau, Natascha Bartlick, Teresa R. Wagner, Theresa Froehlich, Grace Jarjour, Stefan Nueske, Armin Scholz, Bjoern Traenkle, Boris Macek, Ulrich Rothbauer

## Abstract

The mitochondrial outer membrane (MOM)-anchored GTPase Miro1, is a central player in mitochondrial transport and homeostasis. The dysregulation of Miro1 in amyotrophic lateral sclerosis (ALS) and Parkinson’s disease (PD) suggests that Miro1 may be a potential biomarker or drug target in neuronal disorders. However, the molecular functionality of Miro1 under (patho-) physiological conditions is poorly known. For a more comprehensive understanding of the molecular functions of Miro1, we have developed Miro1-specific nanobodies (Nbs) as novel research tools. We identified seven Nbs that bind either the N- or C-terminal GTPase domain of Miro1 and demonstrate their application as research tools for proteomic and imaging approaches. To visualize the dynamics of Miro1 in real time, we selected intracellularly functional Nbs, which we reformatted into chromobodies (Cbs) for time-lapse imaging of Miro1. By genetic fusion to an Fbox domain, these Nbs were further converted into Miro1-specific degrons and applied for targeted degradation of Miro1 in live cells. In summary, this study presents a collection of novel Nbs that serve as a toolkit for advanced biochemical and intracellular studies and modulations of Miro1, thereby contributing to the understanding of the functional role of Miro1 in disease-derived model systems.

## Introduction

Neurodegenerative disorders including Alzheimer’s disease, Parkinsons’s disease (PD) and amyotrophic lateral sclerosis (ALS) pose a major public health challenge especially in societies with a rapidly aging population. Due to the fundamental role of mitochondria in energy production, calcium homeostasis, reactive oxygen species (ROS) formation, and initiation of apoptosis [1], pathological mitochondrial morphologies and a dysfunctional quality control are among the main drivers in the development and progression of neurodegeneration [2, 3]. Within cells, mitochondria are continuously transported along actin- and microtubule-based cytoskeletal pathways to areas in demand of ATP supply or calcium buffering [4, 5]. Microtubule-based transport is effected by the motor proteins kinesins and dyneins in anterograde and retrograde directions, respectively [6]. Miro1, a member of the Rho GTPase family, which is mainly located on the surface of the mitochondrial outer membrane (MOM) [7–9], acts as an adaptor to tether motor complexes to mitochondria. Structurally, Miro1 is composed of two distinct N- and C-terminally located GTPase domains flanking a pair of calcium ion (Ca^2+^) binding EF-hands and a C-terminal transmembrane domain anchored in the MOM [10, 11]. By its cytoplasmic domains, Miro1 interacts with the adaptor proteins TRAK1 and TRAK2 [11], which recruit the motor proteins kinesin-1 (KIF5B) and dynein/dynactin to facilitate mitochondrial transport along microtubules [12, 13]. This motor/adaptor complex is regulated by Ca^2+^ levels. At high concentrations, Ca^2+^ arrests mitochondrial transport by binding to the EF hand domains of Miro1, causing the motor complex to detach from the organelle [14–17]. Similarly, Miro1 interacts with Cenp-F for cell cycle-dependent distribution of mitochondria during cytokinesis [18, 19], and overexpression of Miro1 was shown to enhance intercellular transfer of mitochondria from mesenchymal stem cells to stressed epithelial cells [20, 21]. Besides mitochondria, Miro1 was also shown to play an important role in peroxisomal transport [19, 22].

Additionally, Miro1 is a known substrate of the mitophagy-associated PINK1/Parkin quality control system [23] and impaired Miro1 ubiquitination has been recently linked to Parkin mutants found in Parkinson’s disease (PD) patients and in fibroblasts from an at-risk cohort [24, 25]. This was further confirmed by the identification of mutations in the Miro1 gene *RHOT1* causing an altered mitophagy response [26, 27]. Although not yet demonstrated at the molecular level, dysregulated cellular levels of Miro1 have been described in ALS animal models and in patients [28, 29]. These results, in combination with recent data describing aberrant peroxisomal metabolism in PD patients [27, 30], strongly suggest a multifactorial link between Miro1 and neurological diseases. Consequently, Miro1 is considered as an emerging biomarker and potential drug target in neuropathology [25, 31].

Despite previously conducted *in vitro* and *in vivo* studies on Miro1, detailed information about its structure-related function and cellular dynamics are still lacking [32]. This can be partially attributed to a limited availability of research tools to appropriately study Miro1. Most analyses rely on ectopic expression of fluorescent fusion constructs or epitope-tagged Miro1 [10, 11, 18, 19, 33]. On the other hand, only a limited number of Miro1-specific reagents such as antibodies are available to visualize and study endogenous Miro1.

Single-domain antibody fragments, also known as nanobodies (Nbs), derived from heavy chain-only antibodies found in camelids [34] have been established as attractive alternatives to conventional antibodies for a multitude of biochemical assays [35–38] and advanced imaging applications (reviewed in [39–41]). Additionally, their small size, high solubility and stability qualify Nbs for intracellular expression (reviewed in [42, 43]). Intracellularly functional Nbs genetically fused to fluorescent proteins, designated as chromobodies (Cbs), have been successfully applied for tracing their target antigens in different cellular compartments as well as in whole organisms (reviewed in [39, 44]). For the generation of advanced research tools to study Miro1, we developed specific Nbs from an immunized library. Following an in depth characterization of their binding properties, we selected candidates applicable as capture and detection reagents in biochemical assays, immunofluorescence staining and live-cell imaging. Additionally, we designed specific degrons for selective degradation of Miro1 within living cells. Based on our results, we propose that the presented Nb-based toolkit opens new opportunities for more comprehensive studies of Miro1 and helps to elucidate its multifaceted roles for mitochondrial malfunction in neurological disorders.

## Results

### Identification of Miro1-specific Nbs

To generate Nbs specific for human Miro1 (Miro1), we immunized an alpaca (*Vicugna pacos*) with recombinant Miro1 adopting a 91-day immunization protocol. After confirmation of a successful immune response in a serum ELISA (**Supplementary Figure 1**), we generated a phagemid library of ∼1.5 × 10^7^ clones from peripheral B lymphocytes of the immunized animal, representing the diverse repertoire of variable domains (VHHs or Nbs) of the heavy chain-only antibodies. For selection of Miro1-specific Nbs, we performed phage display using either passively adsorbed Miro1, or GFP-Miro1 from HEK293 cells with site-directed immobilization employing the GFP moiety (**Supplementary Figure 2A**). After two and three rounds of biopanning against Miro1 or GFP-Miro1, we analysed a total of 200 individual clones by phage ELISA and identified 42 positive binders (**Supplementary Figure 2B, C**). Sequence analysis revealed seven unique Nbs with highly diverse complementarity determining regions (CDR) 3 (**Figure 1A**). For further analysis, Miro1-Nbs were produced with a C-terminal His_6_ tag in *Escherichia coli* (*E.coli*) and purified via immobilized metal ion affinity chromatography (IMAC) followed by size exclusion chromatography (SEC) (**Figure 1B**). Binding affinities were assessed by biolayer interferometry (BLI) for which we immobilized biotinylated Nbs on streptavidin (SA) biosensors and measured their binding kinetics by loading different concentrations of Miro1. Our results showed that all seven Nbs bind Miro1 with high affinities in the low nanomolar range with K_D_ values ranging from 1.9 to 29.5 nM (**Figure 1C, Table 1, Supplementary Figure 3A-F**).

**Figure 1.**
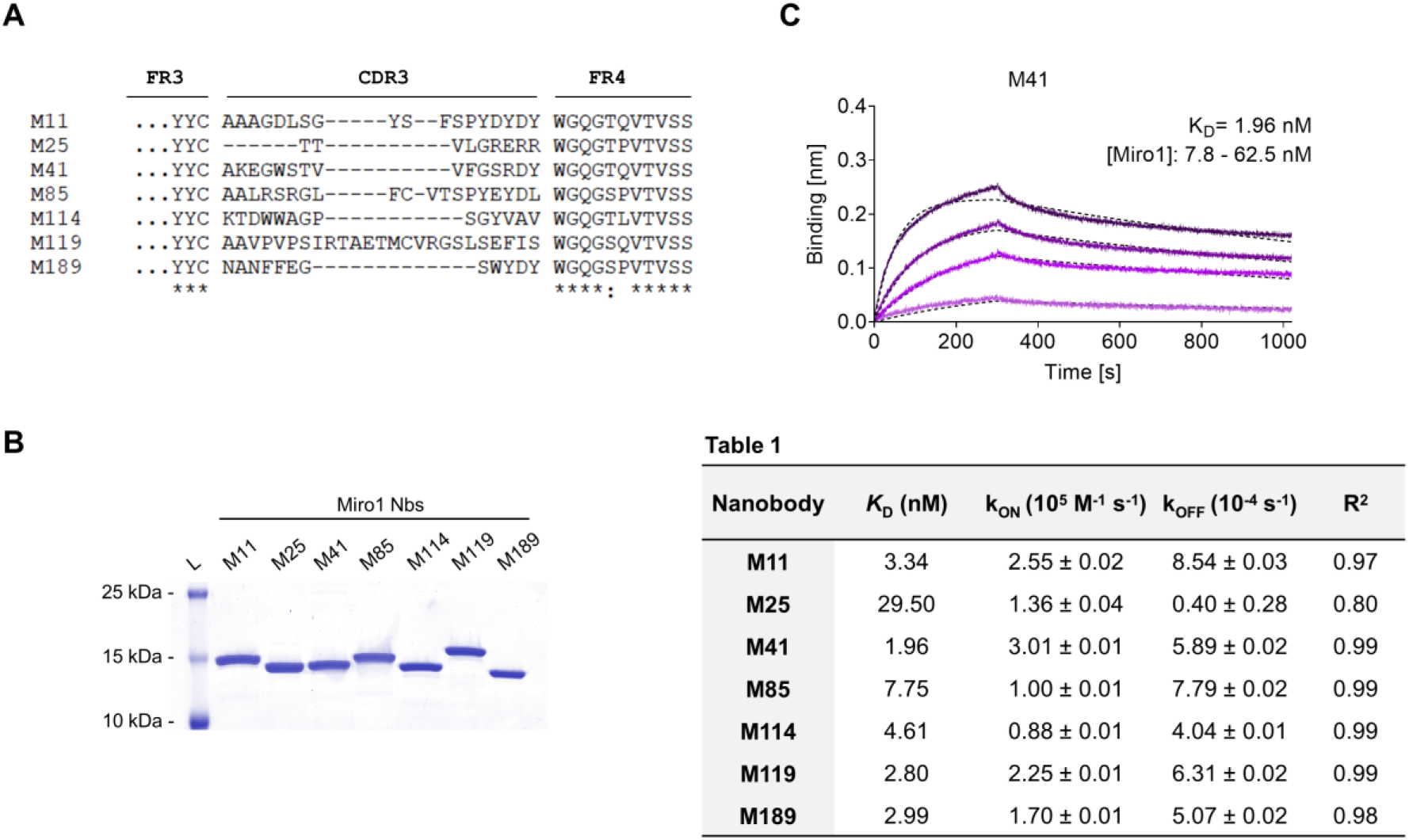
Biochemical characterization of Miro1 specific Nbs. (A) Amino acid sequence alignment of the complementary determining region (CDR) 3 of seven unique Miro1-Nbs positively identified by phage ELISA. (B) Recombinant expression and purification of Nbs using immobilized metal affinity chromatography (IMAC) and size exclusion chromatography (SEC). (C) For biolayer interferometry based affinity measurements, Miro1-Nbs were biotinylated and immobilized on streptavidin sensors. Kinetic measurements were performed by using four concentrations of purified Miro1 ranging from 3.9 nM – 1 µM. As an example, the sensogram of Miro1 on immobilized M41-Nb at indicated concentrations (illustrated with increasingly darker shades from low to high concentration) is shown and global 1:1 fits are illustrated as dashed line (upper panel). The table summarizes affinities (K_D_), association (k_ON_) and dissociation constants (k_OFF_), and coefficient of determination (R^2^) determined for individual Nbs (lower panel).

### Immobilized Miro1-Nbs specifically precipitate Miro1

Considering that a variety of Nbs covalently immobilized to solid matrices such as agarose particles have been applied as pulldown reagents to capture their antigens [35, 45, 46], we used this approach to investigate the functionality of Miro1-Nb candidates for immunoprecipitation (IP). We chemically coupled Miro1-Nbs to N-hydroxysuccinimide (NHS)- activated agarose particles thereby generating Miro1-nanotraps. First, these nanotraps were incubated with the soluble fraction of cell lysates derived from HEK293 cells expressing GFP- Miro1 or GFP as control. Immunoblot analysis of input and bound fractions revealed that all Miro1-nanotraps except M11 and M85 efficiently precipitated GFP-Miro1, with M114 and M119 exhibiting the highest pulldown efficiencies comparable to the commercially available high-affinity GFP nanotrap [35] (ChromoTek) (**Figure 2A**). Notably, none of the tested nanotraps showed unspecific binding to GFP or GAPDH used as an endogenous control (**Figure 2A, Supplementary Figure 4**). Next, we investigated the potential of the nanotraps to precipitate endogenous Miro1 from soluble HEK293 protein extracts. Immunoblot analysis revealed Miro1 in the bound fractions of M41, M114, M119 and M189. The levels of precipitated Miro1 were comparable or slightly higher to those obtained with a conventional anti-Miro1 antibody, while no unspecific binding of Miro1 to a nanotrap displaying a non-related Nb was detected (**Figure 2B**). Because we generally observed rather low levels of endogenous Miro1, we tested different cell types such as U2OS or HeLa cells, different lysis conditions, or the use of different Miro1-specific antibodies for detection (**Supplementary Figure 5**). From these results it can be concluded that only minor fractions of endogenous Miro1 are released under non-denaturing lysis conditions, making it difficult to study Miro1 regardless of the capturing reagent used.

**Figure 2.**
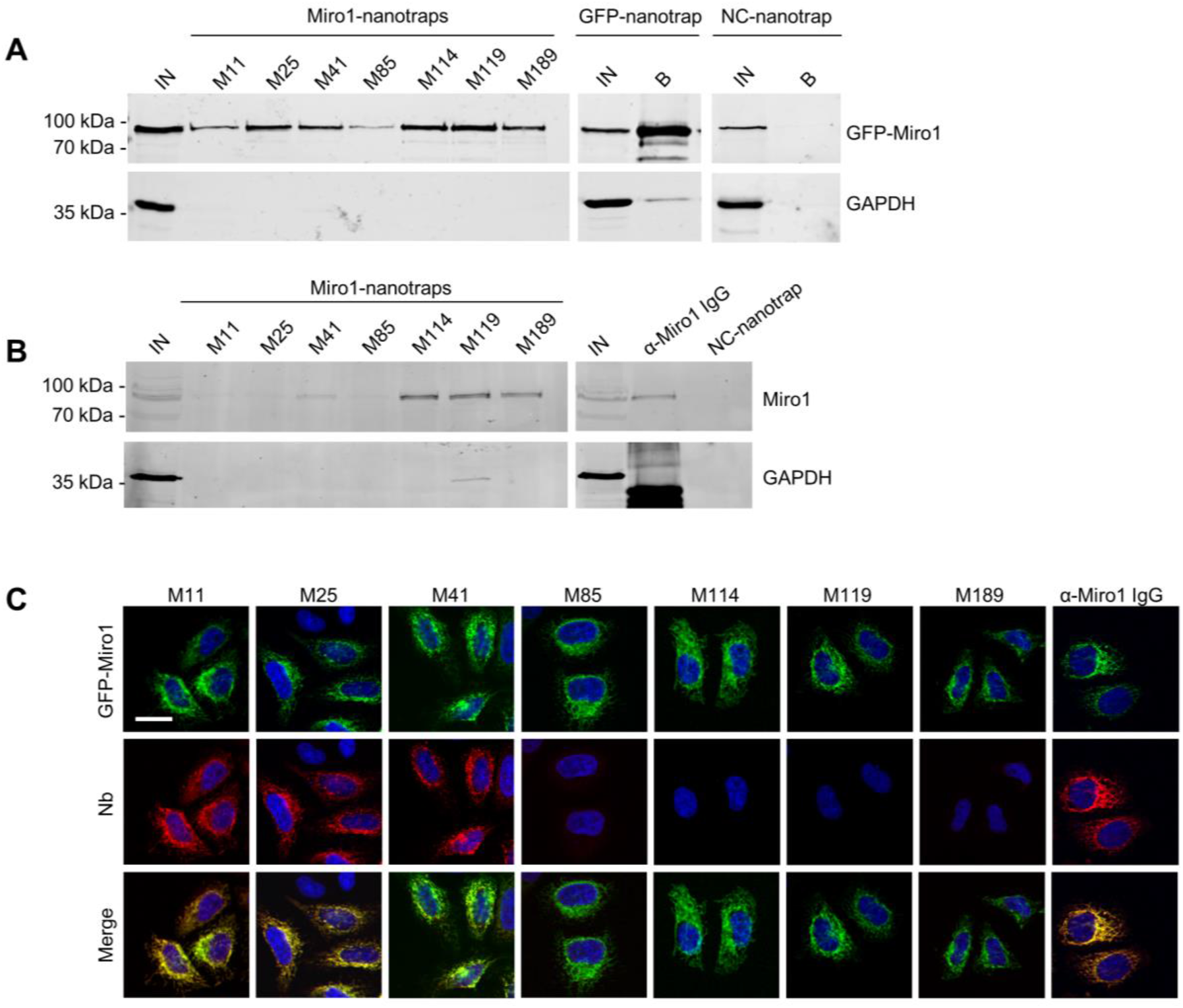
Immunoprecipitation of Miro1 with Nbs. (A) For immunoprecipitation with immobilized Nbs (nanotraps), soluble protein fraction of HEK293 cells transiently expressing GFP-Miro1 or GFP as control, was adjusted to 2 mg/mL and incubated with equal amounts of nanotraps. Input (IN, 1% of total) and bound (20% of total) fractions were subjected to SDS-PAGE followed by immunoblot analysis using antibodies specific for GFP (upper panel) and GAPDH (lower panel). As positive control GFP-nanotrap and as negative a non-specific (NC) nanotrap were used. (B) Immunoprecipitation from non-transfected HEK293 as described in **A** were performed. Input and bound fractions were analysed with an anti-Miro1 antibody. As positive control anti- Miro1 IgG immobilized on Protein A/G sepharose and as negative control a non-specific (NC) nanotrap was used. (C) Immunofluorescence (IF) detection of GFP-Miro1 in fixed and permeabilized HeLa cells after staining with Miro1-Nbs as primary labelling probes. Representative confocal laser scanning (CLSM) images are shown of each individual Nb detected with anti-VHH antibody labelled with Cy5 (middle row). As positive control, transfected cells were stained with anti- Miro1 antibody followed by detection with a Cy5-labelled secondary antibody. Nuclei were counterstained with DAPI. Scale bar 20 µm.

### Immunofluorescence studies with Miro1-Nbs

For a second functional testing, we examined the performance of Miro1-Nbs in immunofluorescence (IF). Therefore we applied the Nbs as primary binding molecules in combination with fluorescently labelled anti-VHH antibodies in fixed and permeabilized HeLa cells transiently expressing GFP-Miro1. Interestingly, Nbs M11 and M25, which did not capture Miro1 by immunoprecipitation showed a clear co-localization with GFP-Miro1 at mitochondrial structures. With M41-Nb, we identified one candidate which displayed functionality in both assay types (**Figure 2C**). Taken together, our data from IP and IF analysis showed the identified Miro1-Nb candidates have the potential to capture and detect their antigen *in vitro*.

### Selected Miro1-Nbs bind different domains of Miro1

To generate well-characterized binders, detailed knowledge of their recognized epitopes or domains is mandatory. Considering that Nbs preferentially bind conformational epitopes [47, 48], we performed domain mapping to identify the binding regions recognized by our Miro1- Nbs. Therefore, we generated a series of Miro1 domain deletion constructs fused C-terminally to GFP (**Figure 3A**) and performed pulldown studies upon expression of these deletion constructs in HEK293 cells as described above. Analysis of the bound fractions revealed that M11, M41, M85, M114 and M189 specifically recognize epitopes within the C-terminal GTPase domain while M25 addresses an epitope spanning the N-terminal GTPase in combination with the EF-hand domains. For M119, we observed interactions involving regions of the EF hands as well as the C-terminal GTPase domains (**Figure 3B**).

**Figure 3.**
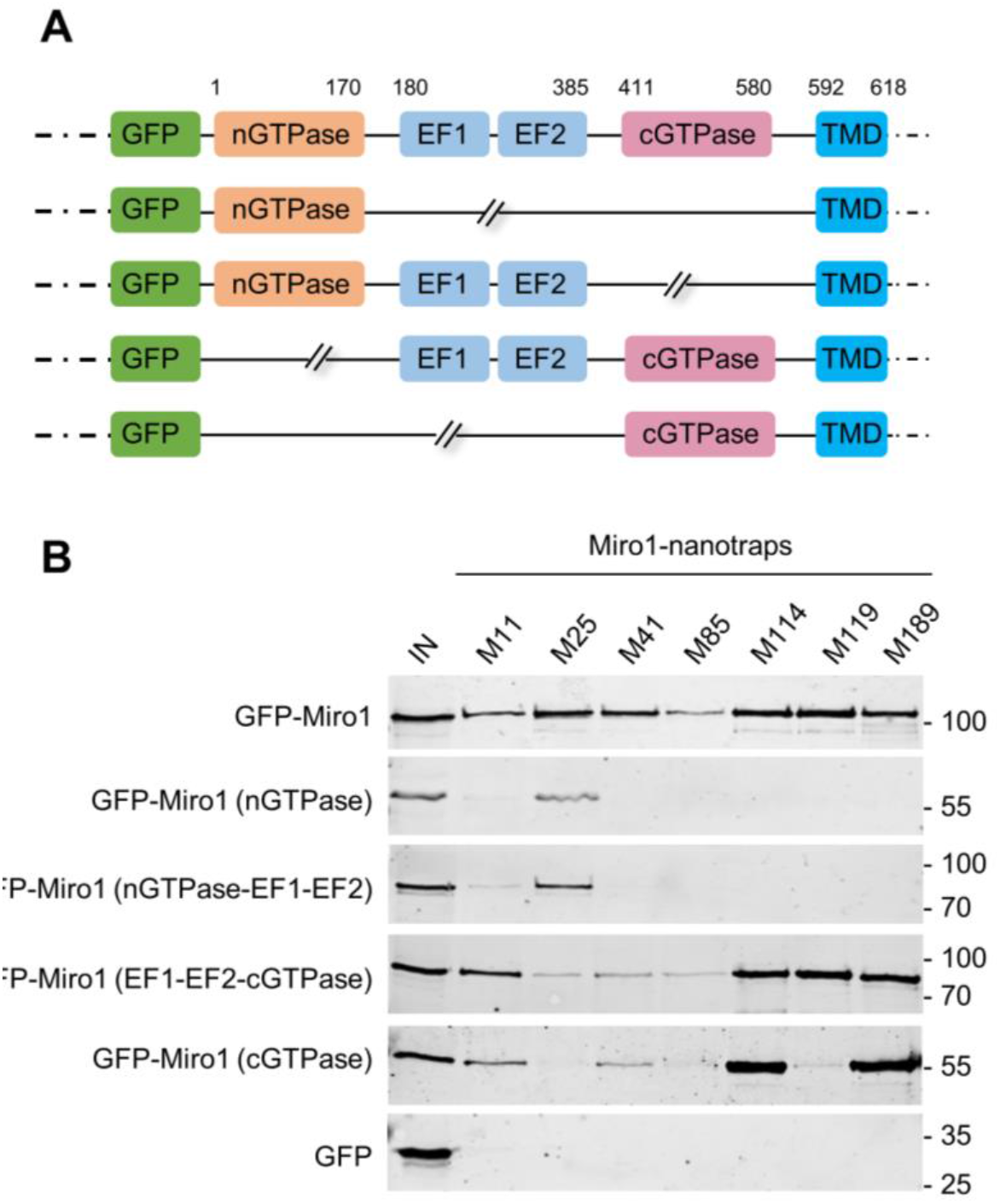
Domain mapping of Miro1-Nbs. (A) Schematic illustration of GFP-labelled Miro1 deletion constructs and domains used for domain specific binding studies. (B) Soluble protein fractions of HEK293 cells transiently expressing indicated Miro1 deletion constructs or GFP (as control) were subjected to immunoprecipitation with selected Miro1 nanotraps followed by western blot analysis of input (IN) and bound fractions with an anti-GFP antibody.

### Optimized, bivalent Nbs show improved capture and detection of Miro1

With M41- and M114-Nb we identified two candidates that have high affinities, are devoid of additional disulfide bonds and tested positive in IP and/ or IF detection of Miro1. In the monovalent format, however, both Nbs bind only small amounts of Miro1. To improve their binding performance, we genetically fused the coding sequences of two M41- or two M114- Nbs head-to-tail, inserting a flexible Gly-Ser linker ((G_4_S)_4_) of 20 amino acids and generated a bivalent M41-Nb (_biv_M41) and a bivalent M114-Nb (_biv_M114). Following production and purification from mammalian cells (**Supplementary Figure 6A**), we analyzed their binding kinetics by BLI measurements showing similar or slightly improved affinities (**Supplementary Figure 6B**). To avoid potential reduction in binding due to nonspecific modification of lysine residues by NHS-based coupling, we changed to a site-specific functionalization strategy. Thus, we introduced an azide-modified peptide at the C-terminus of the bivalent Nbs via chemoenzymatic sortagging [49, 50] followed by click-chemistry addition of a dibenzocyclooctyne (DBCO) derivate. This enabled us to flexibly conjugate either agarose particles or fluorescent dyes specifically to the C-terminus of the bivalent Nbs [51]. With this approach, we converted _biv_M41 and _biv_M114 into fluorescently labeled bivalent imaging probes and nanotraps, which we further tested in their respective applications.

First, we performed IF staining of GFP-Miro1 expressing HeLa cells with either the monovalent Nbs, which were chemically coupled to AlexaFluor (AF) 647 (M41_647_NHS_; M114_647_NHS_) or the bivalent formats, which were C-terminally conjugated to AF647 (_biv_M41_647_sort_; _biv_M114_647_sort_). For _biv_M41_647_sort_, image analysis showed a significantly improved staining and a crisp overlap of the GFP-Miro1 and Nb signal at mitochondrial structures compared to the monovalent version. For M114-Nb, for which no Miro1 was detectable with the monovalent version, mitochondrial structures became visible once this Nb was applied in the bivalent format. Notably, when tested for detection of endogenous Miro1 in HeLa cells, _biv_M114_647_sort_ shows a strong mitochondrial staining comparable to the conventional Miro1 antibody (**Figure 4**).

**Figure 4.**
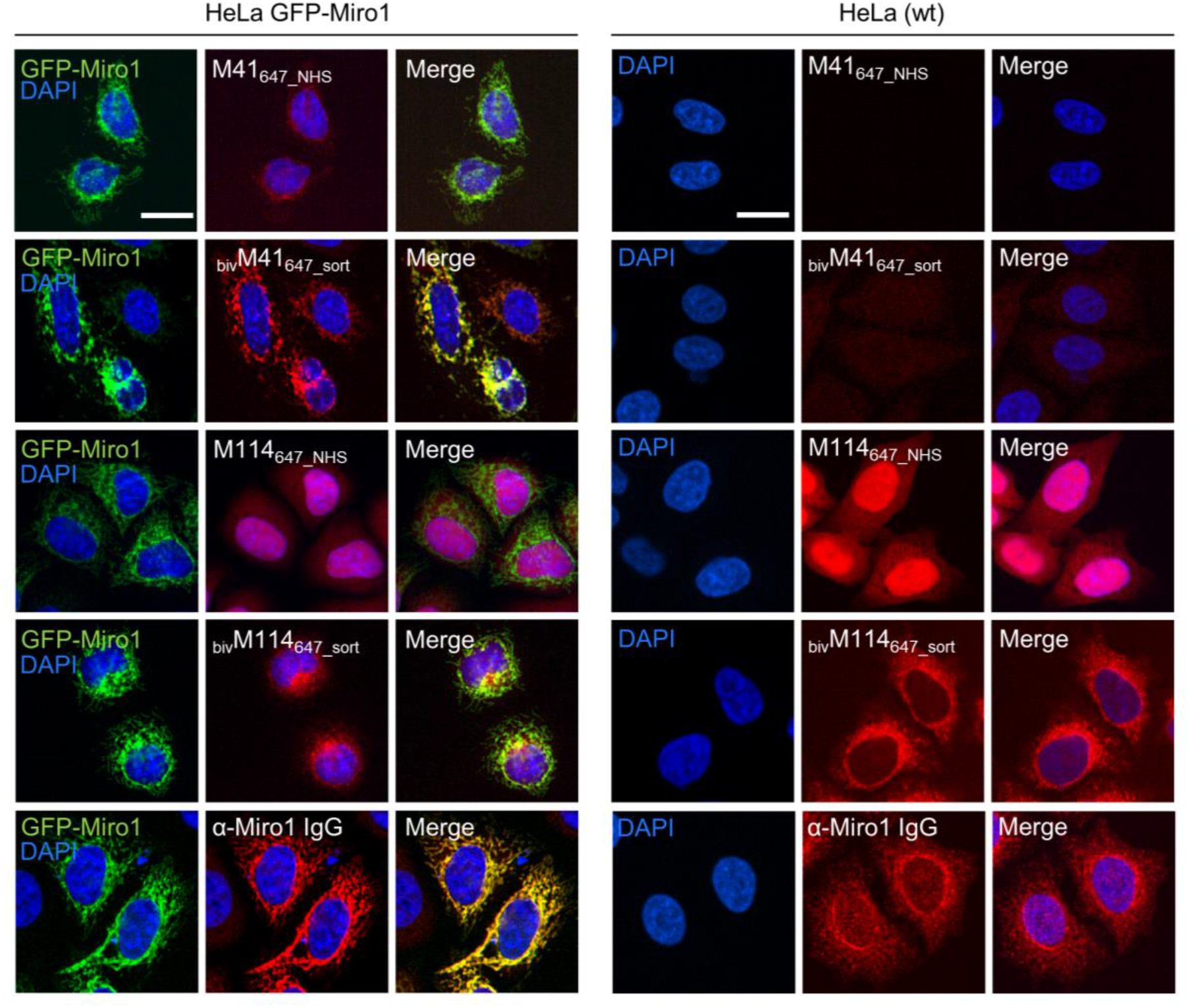
Comparison of monovalent and site specifically conjugated bivalent M41- and M114-Nbs in immunofluorescence. For comparable IF analysis HeLa cell transiently expressing GFP-Miro1 (left panel) or wildtype (wt) HeLa cells (right panel) were fixed and stained with the mono- or bivalent Nbs conjugated either chemically (M41_647_NHS_, M114_647_NHS_) or site-specifically via sortagging (_biv_M41_647_sort_, _biv_M114_647_sort_) to AlexaFluor 647 (647). As positive control respective cells were stained with an anti-Miro1 IgG followed by detection with a secondary antibody conjugated to AlexaFluor 647 (bottom row). Representative fluorescence images are shown from three independent biological replicates. Nuclei were counterstained with DAPI. Scale bar 20 µm.

Next, we tested the different nanotraps to pull down endogenous Miro1. Comparative immunoprecipitation of endogenous Miro1 from lysates of HEK293 cells using either the chemically immobilized monovalent nanotraps (M41_NHS_; M114_NHS_) or the site-directed modified versions (_biv_M41_sort_; _biv_M114_sort_) revealed a considerably increased accumulation of endogenous Miro1 after pulldown with _biv_M41_sort_ and also slightly higher enrichment for _biv_M114_sort_. Notably, in both cases the amount of precipitated Miro1 was higher compared with the conventional Miro1 antibody used as a positive control (**Figure 5A**). For a more detailed analysis, we continued and performed an in-depth comparison of the mono- and bivalent nanotraps (M41_NHS_; M114_NHS_ and _biv_M41_sort_; _biv_M114_sort_) using mass-spectrometry analysis to evaluate their potential use in protein interaction studies. In total, three technical replicates for each nanotrap were performed, with equal cell number as input material, which allowed us to apply label-free quantification. Initially, we validated the reproducibility between replicates, which showed a Spearman rank correlation close to one (**Supplementary Figure 7A**). Correlation was lower between the different monovalent or bivalent nanotraps, suggesting a difference in their performance. These findings were further supported by principal component analysis (PCA), for which we observed that ∼66% of the sample variance can be explained in the first component by the two different nanotrap formats used (**Supplementary Figure 7B**).

**Figure 5.**
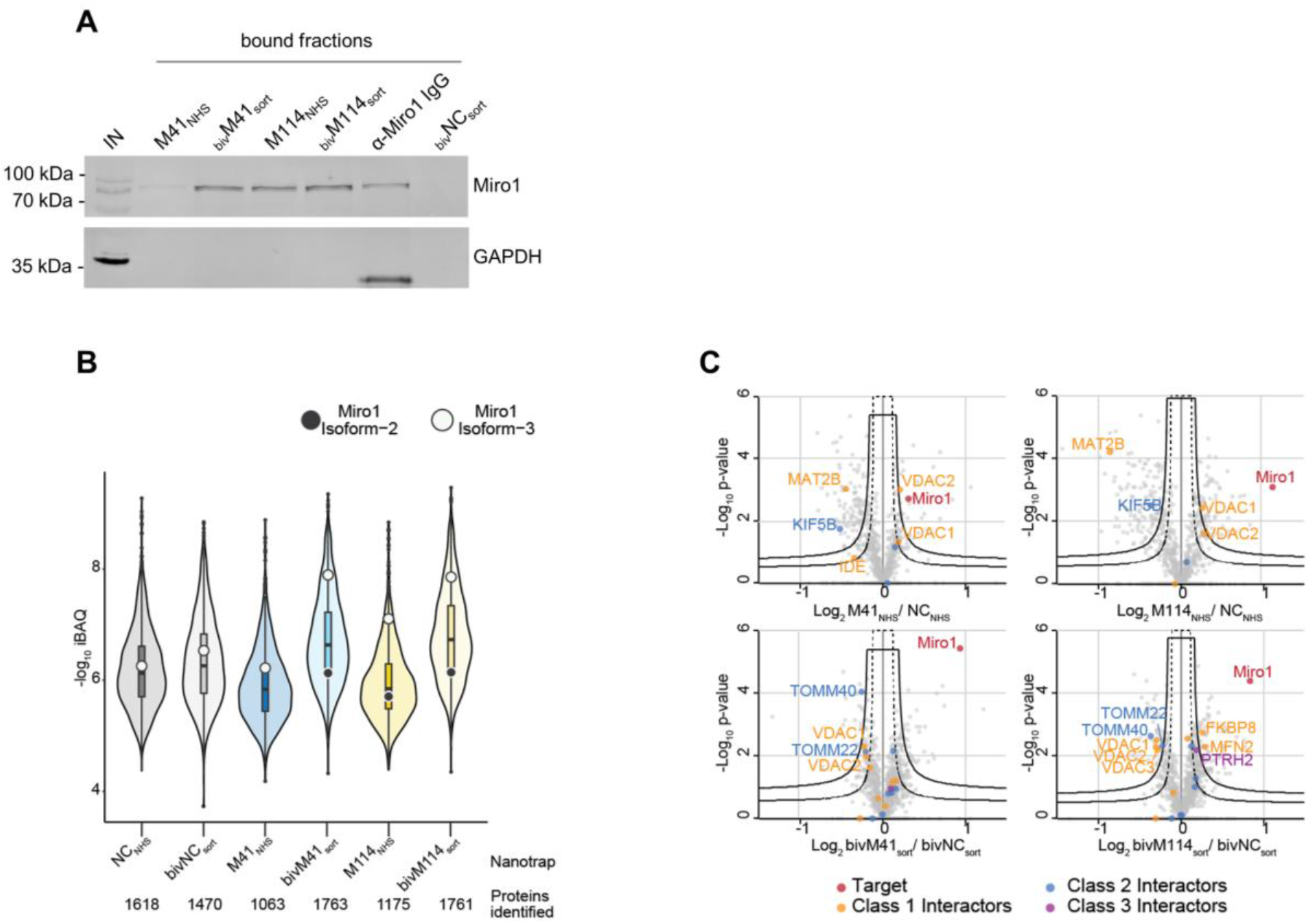
Proteomic analysis of Miro1 capture. (A) For comparable immunoprecipitation soluble protein fraction of HEK293 cells were incubated either with monovalent Nbs either chemically coupled to NHS sepharose (M41_NHS_, M114_NHS_) or the bivalent formats, which were site specifically conjugated to agarose particles by sortagging and click chemistry (bivM41_sort_, bivM114_sort_). Input and bound fractions were analysed with an anti-Miro1 antibody. As positive control anti-Miro1 IgG immobilized on Protein A/G sepharose and as negative control a non-specific bivalent and site specifically conjugated nanotrap (bivNC_sort_) was used. Shown is a representative immunoblot stained with an anti- Miro1 antibody reflecting the results of three independent biological replicates. (B) Capture efficiency by mono- and bivalent nanotraps. Averaged iBAQ (intensity based absolute quantification) values for Miro1 isoform 2 (white circles) and isoform 3 (black circles) of three biological replicates are shown. (C) Classification of Miro1 interactor based on STRING database. Class 1: direct interactor, confidence score >0.9; Class 2: direct interactor, confidence score <0.9; Class 3: indirect interactor, confidence score >0.9.

Next, we analysed the efficiency of each nanotrap to capture endogenous Miro1. Sequence alignment analysis of the eight annotated Miro1 isoforms showed a high level of sequence identity between isoforms (data not shown). In total, 31 “razor” peptides were identified for Miro1 isoform 3 and 30 peptides, including one unique peptide, for isoform 2 (**Supplementary Figure 7C**). Both bivalent nanotraps, as well as the M114_NHS_ nanotrap, were able to capture isoform 2 in addition to isoform 3. Notably, bivalent nanotraps allowed for the identification of more Miro1 peptides compared to their monovalent formats, which is also reflected by the higher Miro1 sequence coverage (**Supplementary Figure 7D**). We validated the enrichment efficiency of Miro1 capture based on intensity based absolute quantification (iBAQ) (**Figure 5B**). Overall, we detected a higher background for bivalent nanotraps of up to 1,760 proteins, which is comparable to the controls. Despite the high background, the highest level of Miro1 isoform 3 and 2 was detected for both bivalent nanotraps. While the M41_NHS_ nanotrap showed comparable levels of Miro1 as the controls, M114_NHS_ as well as the bivalent nanotraps all showed an increased Miro1 capture. Finally, we classified known Miro1 interactors (based on STRING database annotation), by their direct or indirect interaction with Miro1 in combination with the confidence score. While all nanotraps showed a clear enrichment of Miro1, in comparison to the control nanotraps, only the _biv_M114_sort_ allowed for the enrichment of class 1 interactors such as MFN2 or FKBP8 (**Figure 5C**).

In summary, these results demonstrate how the performance of Nbs as capture and detection tools can be improved by generating bivalent binding molecules in combination with site-specific functionalization. From our data we concluded that the bivalent Miro1-Nbs have a high potential as detection probes to visualize even low levels of endogenous Miro1. Similarly, the bivalent nanotraps showed an improved performance in capturing Miro1. Especially the site- specifically modified _biv_M114-Nb might be a suitable capture reagent to be applied in future interactome studies of Miro1.

### Characterization of intracellular binding of Miro1-Cbs

The advantage of Cbs, defined as chimeric expression constructs comprising a Nb genetically fused to a fluorescent protein, is that they can visualize dynamic redistribution and expression levels of endogenous antigens within living cells with spatial and temporal resolution (reviewed in [44]). To analyse the functionality of selected Miro1-Nbs for intracellular targeting and tracing of Miro1 in living cells, we converted them into Cbs by fusing the Nb-coding sequences via a flexible GS linker to TagRFP. The Cb constructs were transiently expressed either alone or in combination with GFP-Miro1 in HeLa cells followed by live-cell fluorescence imaging. For M41-, M85- and M114-Cb, we observed a clear relocalization of the Cb signal to mitochondrial networks in the presence of GFP-Miro1 (**Figure 6A**), whereas all other Miro1-Cbs seem to lose their binding properties or could not access their epitopes within the cellular localized antigen (**Supplementary Figure 8**). Additionally, we examined whether the intracellular functional M41-, M85- and M114-Cb could recognize the previously identified domains of Miro1 within living cells. Thus, we expressed the Miro1 domain deletion constructs described above (**Figure 3A**) along with the Cb constructs in HeLa cells. Subsequent cellular imaging showed that ectopic expression of the C-terminal GTPase domain of Miro1 resulted in specific relocalization of M41-, M85- and M114-Cb to the mitochondrial network, confirming our results from pulldown domain mapping for these Cbs (**Supplementary Figure 9A-C)**. However, when we examined Cb binding to endogenous Miro1, we observed a rather diffuse cellular distribution of the Cb signal and we did not detect any characteristic mitochondrial structures as seen with an anti- Miro1 antibody staining of Cb expressing cells (**Supplementary Figure 9D**).

**Figure 6.**
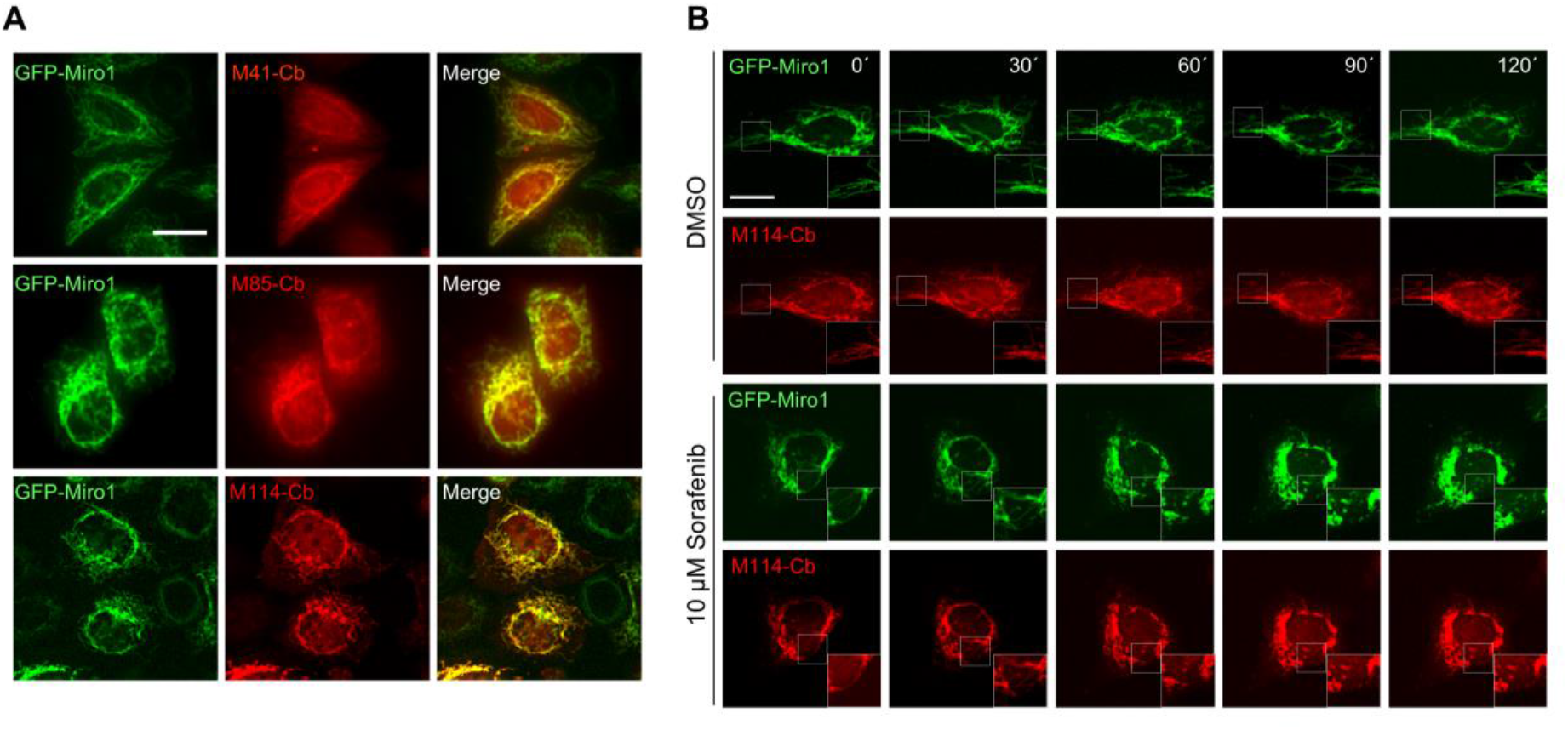
Live-cell imaging of Miro1 with selected Miro1-Cbs. (A) Representative fluorescence images of living HeLa cells transiently expressing GFP-Miro1 (left column) in combination with red fluorescently labelled (TagRFP) M41-, M85- or M114-Cb (middle column). Scale bars 20 µm. (B) Time-lapse microscopy of U2OS cells transiently expressing GFP-Miro1 in combination with either M114-Cb or mitoMkate2. To visually track morphological mitochondrial changes, cells were treated with either DMSO as a control (top two rows) or 10 µM Sorafenib (bottom two rows) followed by time-lapse imaging over a 2 hour period. Shown are representative images of three biological replicates. Scale bar 25 µm. Squares at the bottom right represent enlargements of the selected image section.

### Visualization of compound-induced mitochondrial dynamics in living cells

Our Cb-based imaging results indicated that the M114-Cb shows a slightly better intracellular binding compared to M41- and M85-Cb. Thus, we continued and investigated the utility of this Cb as an intracellular biosensor to track changes in mitochondrial morphology in living cells. For real-time analysis, U2OS cells transiently co-expressing the M114-Cb or a mito-mKate2 construct and GFP-Miro1 were treated with Sorafenib or DMSO as control. Sorafenib has been shown to induce mitochondrial fragmentation and apoptosis in a time-dependent manner [52, 53]. By time-lapse imaging, we visualized changes in mitochondrial morphology within single cells over a two hour period with an imaging interval of 30 min (**Figure 6B, Supplementary Figure 10**). Following 30 min of Sorafenib treatment, images revealed the collapse of the mitochondrial network reflected by gradual disappearance of elongated mitochondria. After 90 min, condensed mitochondrial networks were visible in the majority of treated cells. Both, the Sorafenib induced mitochondrial morphological transitions as well as normal shape changes observable in the DMSO control were successfully visualized by the M114-Cb. These results underline the applicability of the M114-Cb to monitor morphological changes of mitochondria in real time.

### Selective degradation of Miro1 in living cells

Previously, it was reported that depletion of Miro1 by pharmacological intervention or genetic silencing rescues mitophagy activation [31]. To test whether our Nbs could also be used to induce targeted degradation of Miro1 in living cells, we engineered genetically encoded Miro1- specific degrons. To this end, we fused the intracellularly functional Nbs, M41 and M114, or the GFP-Nb (GBP) as control, N-terminally to the mammalian Fbox domain to generate a specific loading platform at Miro1 for components of the mammalian E3 ligase complex, namely SKP1 and Cul1 (**Figure 7A**). Notably, similar approaches using other Nbs were successfully applied to induce selective protein knockdown within cells or organisms [54–58]. These Fbox-Nb constructs were cloned into mammalian expression vectors containing an independently transcribed nuclear TagRFP (TagRFP-NLS) to facilitate the identification of transfected cells in cellular imaging analysis. First, we examined whether Fbox-GBP, Fbox- M41, and Fbox-M114 are functional binders and can bind transiently co-expressed GFP-Miro1 in HeLa cells. A clear overlap of the Fbox-Nb signal with GFP-Miro1 after IF staining using an anti-VHH antibody showed that N-terminal fusion of the Fbox domain did not affect intracellular binding of the Nbs (**Supplementary Figure 11A**). Furthermore, quantification of fluorescence intensity in cells coexpressing the Fbox-Nb constructs and GFP-Miro1 revealed a ∼80% reduction of the GFP signal in cells expressing Fbox-M114, ∼50% in cells expressing Fbox- M41, and ∼50% in cells expressing Fbox-GBP, respectively. Co-expression of a nonspecific Nb-Fbox construct (Fbox-NR) results only in a non-significant reduction in GFP-Miro1 (**Supplementary Figure 11B-C**). Although we could not detect a clear relocalization of intracellularly expressed Nbs to endogenous Miro1 before, we continued and investigated the Fbox fusions with respect to their potential to degrade endogenous Miro1. Therefore, HeLa cells were transfected with Fbox-Nbs and cellular levels of Miro1 following 24 h of expression were monitored by quantitative IF imaging using an anti-Miro1 antibody in cells displaying a nuclear TagRFP signal. While characteristic mitochondrial structures were still observable (**Figure 7B**), quantification of the antibody signals in a statistically relevant number of cells (n>500 cells) showed a ∼16% and ∼30% decrease in the IF signal upon expression of Fbox- M41 and Fbox-M114, respectively. Notably, expression of the nonspecific Fbox-NR results only in a minor reduction of less than ∼4% of the Miro1 signal (**Figure 7C**). From these results we concluded that both specific Fbox-Nb fusion constructs can address endogenous Miro1 in live cells and induce targeted degradation of their antigen.

**Figure 7.**
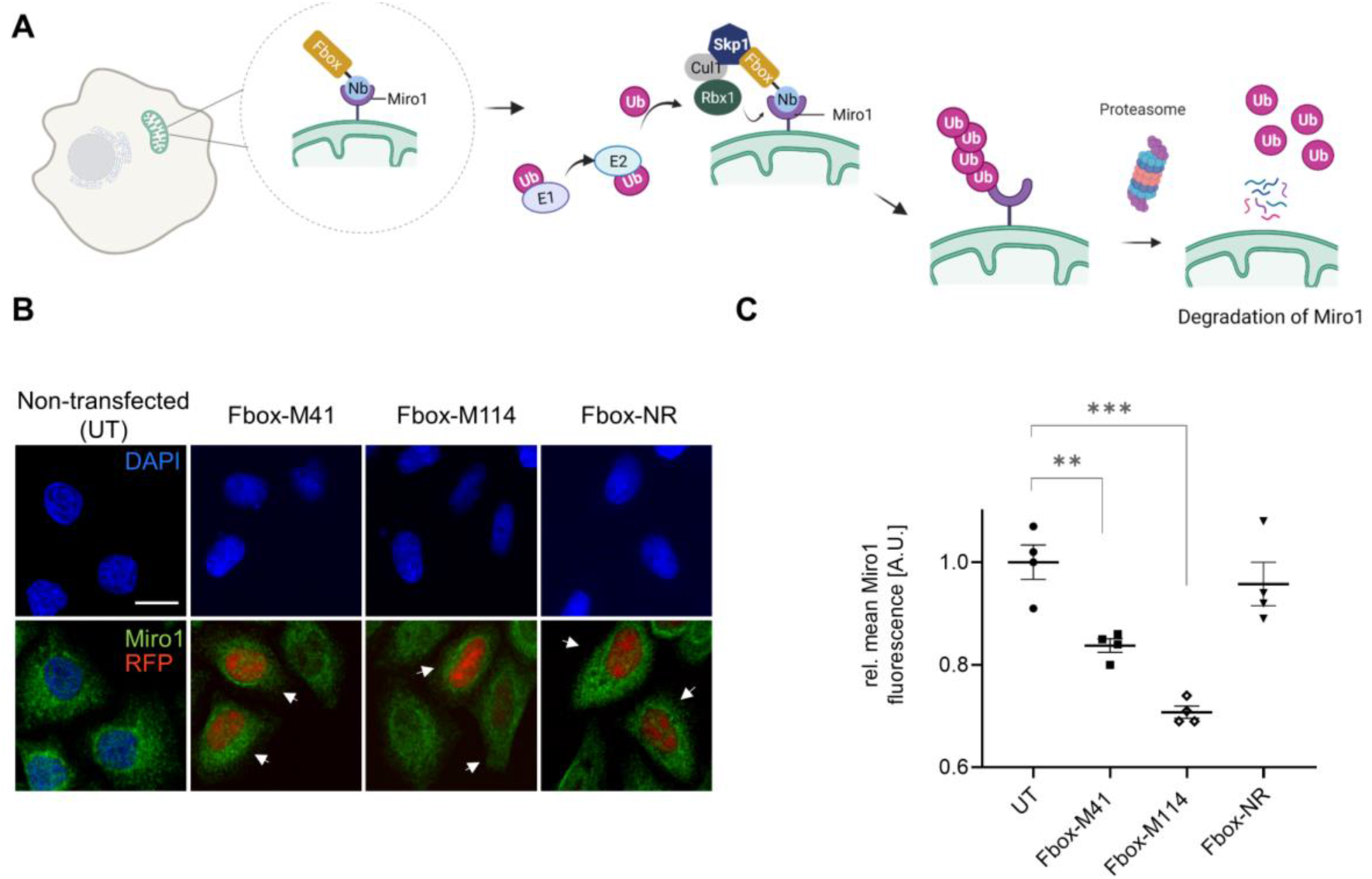
Targeted intracellular degradation of endogenous Miro1 by Fbox-Nb-based degrons. (A) Schematic illustration of the targeted degradation of Miro1 mediated by Miro1-Nb-Fbox fusions (illustration created with Biorender.com) (B) Representative confocal images of HeLa cells transiently expressing indicated Miro1- specific Fbox-Nbs (Fbox-M41, Fbox-M114) or a non-related Fbox-Nb (Fbox-NR) construct. For quantitative IF analysis, cells were fixed and permeabilized 24 h after transfection followed by detection of endogenous Miro1 with Miro1 antibody (shown in green). Fbox-Nb expressing cells were identified by a nuclear TagRFP signal indicated by white arrows and subjected to automated image analysis and quantification as describe in Material and Methods. Scale bar 20 µm. (C) Mean Miro1 fluorescence intensity from HeLa cells expressing Fbox-Nbs determined by quantitative fluorescence imaging. Mean Miro1 fluorescence was calculated from four samples (n= 4; >500 cells) and normalized to untransfected cells, UT (set to 1). As control, a non-related Fbox-Nb construct (Fbox-NR) was used. Data are represented as mean ± SEM. For statistical analysis, student‘s t-test was performed, **p < 0.01, ***p < 0.001.

## Discussion

The emerging role of Miro1 in the development and progression of diseases, particularly neurological disorders, underscores the need for new methods and tools to study this protein in detail at the molecular level [25, 59–62]. Currently, most studies rely on expression of exogenous tagged Miro1 [18, 22, 62, 63]. However, this has been shown to affect mitochondrial morphology and transport in living cells [11] and can also lead to biases in proteomic data, for example, to elucidate potential interaction partners. To expand the ability to study Miro1 in different experimental settings, we developed a collection of Nbs, which were screened for their performance as i) affinity capture tools, ii) labelling probes for fluorescence imaging, iii) intrabodies for visualization and monitoring of Miro1 in live cells, and iv) as Miro1-specific degrons. In total, we selected seven specific binding molecules, which recognize distinct domains of Miro1 with affinities down to the low nanomolar range. These Nbs can be easily produced in high yields in bacteria and four of them were successfully functionalized by simple chemical conjugation as capture molecules to precipitate Miro1 from soluble cell lysates in the monovalent format.

After determining that the identified Nbs in their monovalent format were not suitable as primary probes for visualization of endogenous Miro1, we decided to convert the most promising candidates, M41 and M114, to a bivalent format. In combination with an advanced labeling strategy using site-specific functionalization via sortase tagging in combination with click chemistry, we were able to generate highly functional capture reagents. Our mass spectrometry data indicate that these modified bivalent Miro1 nanotraps are well suited to capture different isoforms of Miro1 and thus have high potential for future interactome studies. Interestingly, only the site-directed modified bivalent M114-Nb also allowed detection of endogenous Miro1 by immunofluorescence imaging. Consistent with previous findings, this confirms the critical impact of Nb formatting and functionalization for the generation of efficient binding molecules that enable one-step detection of their antigens [50, 64, 65]. Considering that IF staining of Miro1 with these bivalent Nbs does not require a secondary antibody and thus a greater spatial proximity of the fluorophore to the target can be achieved, we assume that especially with the _biv_M114-Nb, which also strongly recognizes endogenous Miro1, a promising candidate for the visualization of Miro1 has been generated. We have already developed and used such Nb-based imaging probes for the detection of endogenous vimentin using stochastic optical reconstruction microscopy (STORM) [39, 50]. Thus, we anticipate that the _biv_M114 can be similarly adapted for such advanced imaging techniques on more relevant cells including primary neurons or neurons derived from induced pluripotent stem cells of PD patients which might offer an unprecedented insight in the cellular localization of Miro1 in those models.

Besides their application as recombinant capturing and detection tools, numerous studies reported how Nbs can be functionally expressed in the reducing milieu of living cells to visualize subcellular antigen location or to modulate their target structure and function [43, 45, 66–68]. Accordingly, with M41, M85 and M114, we identified three Nbs, which we converted into intracellularly functional Cbs to visualize Miro1 in living cells. Although, we were not able to detect endogenous Miro1 with Cbs probably due to low levels and a disperse localization of different endogenous isoforms of Miro1, these Cbs could visualize dynamic changes of exogenous Miro1 in time-lapse imaging series. From our treatment studies with Sorafenib we concluded that the Cbs are unaffected by changes in the mitochondrial membrane potential and stably bind their antigen after fixation. This might be advantageous compared to other fluorescent dyes e.g. the MitoTracker series, which bind to thiol groups within mitochondria and have been shown to be sensitive to changes in the membrane potential [69].

To mimic Parkin-mediated Miro1 degradation in living cells, we decided for the application of an artificial nanobody-coupled ubiquitin ligase system. Therefore, we used the M41-Nb and M114-Nb as specific substrate recognition modules for the SKP1-Cul1-Fbox complex to mediate the selective degradation of endogenous and exogenous Miro1. It has to be noted that expression of these degrons did not result in complete depletion of endogenous Miro1. It is possible that the efficacy is due to insufficient binding of endogenous Miro1 yet, which could be further investigated e.g. by FRAP (fluorescent recovery after photobleaching) experiments [45, 70]. Alternatively, it can be speculated whether other known degron entities, such as fusion with TRIM21 [71] or the auxin-dependent degradation system [72], might be better suited for depletion of Miro1 using the target-specific Nbs identified here. Although expression of the degron did not completely deplete endogenous Miro1, the effect was replicable and not observed in the absence of the Fbox protein or in the presence of an unrelated Nb-based degron. For functional studies, CRISPR and RNAi approaches are currently employed to knock down endogenous Miro1 in loss-of-function studies. However, apart from the inherent limitations of each approach, both methods lead to complete loss of Miro1 disrupting mitochondrial homeostasis [59, 73]. For this reason, the Nb-based degrons holds great benefit as an option for targeted Miro1 depletion. With the ongoing development of an inducible system applied in PD neuronal models, Fbox-Nb-mediated Miro1 degradation could expand the possibilities to study PD-related mitophagy impairment.

In summary, this study introduces for the first time, an adaptable and flexible toolset of specific Nbs for the multi-faceted study of the mitochondrial GTPase, Miro1. In the nanotrap format, the identified Nbs are promising capture tools for the proteomic characterization of Miro1 domain-dependent interactions. Conjugated to fluorophores, Nbs have a distinct potential to be applied as labelling probes for one step detection e.g. in high-resolution imaging of Miro1 at mitochondria. Formatted into Cbs, formidable intracellular biosensors of Miro1 dynamics within live cells were generated. As intracellular Miro1 binding agents, the Nbs can be easily fused with different protein domains for the targeted modulation of Miro1 in live cells. The versatile applicability of our Nb set underscores its substantial potential as Miro1-specific research toolkit, and we propose that its scope will soon expand to decipher novel and/or further confirm proposed functions of Miro1 in pathophysiological relevant states.

## Materials and Methods

### Nanobody library construction

Alpaca immunization with purified Miro1 and Nb library construction were carried out as previously described [66]. Animal immunization was approved by the government of Upper Bavaria (Permit number: 55.2-1-54-2532.0-80-14). In brief, one alpaca (*Vicugna pacos*) was immunized with recombinant human Miro1 expressed in *E.coli.* After an initial priming dose of 1 mg, the animal received booster injections of 0.5 mg after the 3^rd^, 4^th^, 7^th^ and 12^th^ week. 20 mL serum samples collected after the 9^th^ and 13^th^ week were analysed for seroconversion. 13 weeks after the initial immunization, 100 mL of blood was collected and lymphocytes were isolated by Ficoll gradient centrifugation using the Lymphocyte Separation Medium (PAA Laboratories GmbH). Total RNA was extracted using TRIzol (Life Technologies) and mRNA was reverse transcribed to cDNA using a First-Strand cDNA Synthesis Kit (GE Healthcare). The Nb repertoire was isolated in 3 nested PCR reactions using following primer combinations: (i*) CALL001* and *CALL002*, (ii) forward primer set *FR1-1*, *FR1-2*, *FR1-3*, *FR1-4* and reverse primer *CALL002*, and (iii) forward primer *FR1-ext1* and *FR1-ext2* and reverse primer set *FR4- 1*, *FR4-2*, *FR4-3*, *FR4-4*, *FR4-5* and *FR4-6* introducing *SfiI* and *NotI* restriction sites. The sequences of all primers used in this study are shown in **Supplementary Table 1**. The Nb library was subcloned into the *SfiI/ NotI* sites of the pHEN4 phagemid vector [74].

### Nanobody screening

For the selection of Miro1-specific Nbs, two and three consecutive phage enrichment rounds were performed either with recombinant Miro1 immobilized on Nunc^TM^ Immuno^TM^ MaxiSorp^TM^ tubes (Thermo Scientific) or HEK293-expressed GFP-Miro1 immobilized on ChromoTek GFP- Trap^®^ Multiwell plate (Proteintech). To generate Nb-presenting phages, *E.coli* TG1 cells comprising the Miro1-Nb library in pHEN4 vector were infected with the M13K07 helper phage. . 1 x 10^11^ phages, prepared from culture supernatant by PEG precipitation, were used for each panning process. Extensive blocking of antigen and phages was performed in each selection round with 5% milk or BSA in PBST (PBS, 0.05% Tween 20, pH 7.4) [48].

For the selection process using recombinant Miro1, phages were first applied on immunotubes coated with GFP (10 µg/mL) to deplete non-specific binders and then transferred to immunotubes either coated with Miro1 (10 µg/mL) or GFP (10 µg/mL) as non-related antigen. Incubation steps were performed at RT for 2 h. Washing stringency was increased for each selection round. Bound phages were eluted in 100 mM triethylamine (pH 12.0), followed by immediate neutralization with 1 M Tris/HCl pH 7.4. For the panning process using GFP-Miro1, 2 × 10^7^ HEK293 cells transiently expressing GFP-Miro1 or GFP were harvested and lysed as previously described [75]. GFP-Miro1 and GFP were immobilized respectively on GFP-Trap^®^ Multiwell plates according to manufacturer’s protocol. To deplete GFP-specific binding molecules, phages were first applied into wells displaying GFP and then transferred into wells with immobilized GFP-Miro1. All incubation steps of three consecutive selection rounds were performed at 4 °C for 2 h under constant mixing. Washing and elution steps were carried out equally as described above. After each panning round exponentially growing *E.coli* TG1 cells were infected with eluted phages and spread on selection plates to rescue phagemids. Antigen-specific enrichment for each round was monitored by counting colony forming unit (CFUs). Following panning, 100 individual clones from both panning strategies were screened by phage ELISA procedures using immobilized Miro1 or GFP-Miro1 as antigen and GFP as control. Bound phages were detected using a horseradish peroxidase-labeled anti-M13 monoclonal antibody (GE Healthcare) and Thermo Scientific^TM^ 1-Step^TM^ Ultra TMB solution.

### Expression Plasmids

For the bacterial expression, Nb sequences were cloned into the pHEN6C vector [35], thereby adding a C-terminal sortase tag (LPETG) followed by a His_6_ tag for IMAC purification as described previously [50]. For mammalian expression, the coding sequence for _biv_M41 was produced by gene synthesis (Thermo Fisher Scientific) and cloned into the pCDNA3.4 expression vector downstream of an N-terminal signal peptide (MGWTLVFLFLLSVTAGVHS) for secretion using restriction enzymes *XbaI* and *AgeI* sites. _biv_M114 was generated by insertion of two coding sequences of M114 into the pCDNA3.4 expression vector in three steps: first, M114 sequence was amplified in two separate PCRs using primer sets *bivM114GA-for*, *nterm1273_rev* and *bivM114GA2_for*, *downEcoRI_rev*; second, both amplicons were then fused by overlap-extension PCR with additional use of primers *bivM114FPCR_for* and *bivM114FPCR_rev*; third, the resulting sequence was cloned into *Esp3I*- and *EcoRI-* digested pCDNA3.4 expression vector by Gibson assembly according to the manufacturer’s protocol. To generate Miro1-Cbs, Nb sequences were genetically fused with TagRFP by ligation into *BglII*- and *BstEII*-sitesof the plasmid previously described as PCNA-chromobody [70]. The expression construct for GFP-Miro1 was generated by Gibson assembly cloning [76] of three fragments: the pEGFP-N1 vector (Takara Bio) backbone, amplified with the primer set *vectorGA_for* and *vectorGA_rev* and the cDNA of human Miro1 isoform 1 (UniProtKB Q8IXI2- 1), amplified in two fragments from the Miro1-V5-HisA plasmid, a gift from Julia Fitzgerald [26] using the primer set *Miro1fragA_for* and *Miro1fragA_rev* and primer set *Miro1fragB_for* and *Miro1fragB_rev* respectively. Miro1 domain deletion constructs for mammalian expression were cloned from the GFP-Miro1 expression plasmid generated in this study. Respective domains and vector backbone were amplified by PCR using the following primer sets: *nGTP_for* and *nGTP_rev* for eGFP-nGTPase-TMD; *nGTP_for* and *nGTPEF2_rev* for eGFP- nGTPase-EF1-EF2-TMD, *ΔnGTP_for* and *ΔnGTP_rev* for eGFP-EF1-EF2-TMD. Amplicons comprising additional terminal KpnI recognition sites were purified, digested with *KpnI/DpnI* and intramolecular re-ligated according to standard protocols. The eGFP-cGTPase-TMD expression plasmid was generated by site directed mutagenesis with the primers, *cGTP_for* and *cGTP_rev* using the Q5 Site-Directed Mutagenesis Kit (New England Biolabs) according to the manufactureŕs protocol. The mammalian expression construct for GFP was previously described [46]. For molecular cloning of the mammalian expression vector pcDNA3_Fbox-Nb-IRES-tRFP-NLS, DNA assembly of the following three fragments was performed. Fragment 1 - the complete sequence of plasmid pcDNA3_NSlmb-vhhGFP4, a gift from Markus Affolter (Addgene plasmid #35579) [56] amplified by PCR using primers *NM95_for* and *NM95_rev*; fragment 2- theIRES sequence, amplified by PCR from a pcDNA3.1 vector variant, pcDNA3.1(+)IRES-GFP, a gift from Kathleen_L Collins (Addgene plasmid #51406) with primers *frag2IRES_for* and *frag2IRES_rev and* fragment 3, generated in two steps. First, an NLS sequence was inserted downstream of the TagRFP sequence in the Cb expression vector described above with primers *nls-insert_for* and *nls-insert_rev* using the Q5 Site-Directed Mutagenesis Kit (NEB) and the resulting plasmid was used as a template to subsequently amplify the TagRFP-NLS sequence using the primers *frag3-tRFP-nls_for* and *frag3-tRFP-nls_rev*. Fragment assembly was carried out using NEBuilder HiFi DNA assembly Master Mix (New England Biolabs) according to manufactureŕs protocol. In the resulting pcDNA3_Fbox-Nb-IRES-tRFP-NLS plasmid the GFP-Nb (vhhGFP4) was replaced by Miro1- Nbs using BamHI and BstEII restriction sites. All generated expression constructs were sequence analyzed after cloning. For fluorescent labeling of mitochondria in living cells, the mammalian expression vector pmKate2-mito (Evrogen plasmid #FP187) was used.

### Protein expression and Purification

Miro1-Nbs were expressed and purified as previously described [44, 45]. Bivalent Nbs were expressed using the ExpiCHO™ system (Thermo Fisher Scientific) according to the manufacturer’s protocol. For quality control, all purified proteins were analyzed by SDS-PAGE according to standard procedures. Protein samples were denatured (5 min, 95 °C) in 2x SDS-sample buffer containing 60 mM Tris/HCl, pH 6.8; 2% (w/v) SDS; 5% (v/v) 2-mercaptoethanol, 10% (v/v) glycerol, 0.02% bromphenol blue prior to analysis. All proteins were visualized by InstantBlue Coomassie (Expedeon) staining. For immunoblotting, proteins were transferred to nitrocellulose membrane (GE Healthcare) and detection was performed using anti-His primary antibody (Penta-His Antibody, #34660, Qiagen) followed by donkey-anti-mouse secondary antibody labeled with AlexaFluor647 (Invitrogen). A Typhoon Trio scanner (GE-Healthcare, excitation 633 nm, emission filter settings 670 nm BP 30) was used for the readout of fluorescence signals.

### Affinity measurements by biolayer interferometry (BLI)

Analysis of binding kinetics of Miro1-specific Nbs was performed using the Octet RED96e system (Sartorius) according to the manufacturer’s recommendations. In brief, 2 - 10 µg/mL solution of biotinylated Miro1-Nbs diluted in Octet buffer (HEPES, 0.1% BSA) was used for 40 s to immobilize the Nb on streptavidin coated biosensor tips (SA, Sartorius). In the association step, a dilution series of Miro1 ranging from 3.9 nM - 1 µM were reacted for 300 s followed by dissociation in Octet buffer for 720 s. Every run was normalized to a reference run using Octet buffer for association. Data were analyzed using the Octet Data Analysis HT 12.0 software applying the 1:1 ligand-binding model and global fitting.

### Cell culture, transfections and compound treatment

HEK293 and U2OS cells were obtained from ATCC (CRL3216, HTB-96), HeLa Kyoto cell line, (Cellosaurus no. CVCL_1922) was obtained from S. Narumiya (Kyoto University, Japan) and HAP1 cells were obtained from Horizon Discovery (UK, catalog number C631). The cell lines were tested for mycoplasma using the PCR mycoplasma kit Venor GeM Classic (Minerva Biolabs, Berlin, Germany) and the Taq DNA polymerase (Minerva Biolabs). Since this study does not include cell line-specific analysis, cell lines were used without additional authentication. Cell lines were cultured according to standard protocols. Briefly, growth media containing DMEM (high glucose, pyruvate, with GlutaMAX™, Thermo Fisher Scientific) supplemented with 10% (v/v) fetal calf serum (FCS, Thermo Fisher Scientific) and 1% (v/v) penicillin/streptomycin (Thermo Fisher Scientific) was used for cultivation. Cells were routinely passaged using 0.05% trypsin–EDTA (Thermo Fisher Scientific) and were cultivated at 37 °C in a humidified chamber with a 5% CO_2_ atmosphere. Transient transfection of U2OS and HeLa Kyoto cells with Lipofectamine 2000 (Thermo Fisher Scientific) was carried out according to manufacturer’s instruction. HEK293 cells were transfected with Polyethyleneimine (PEI, Sigma Aldrich) as previously described [46, 77].Compound treatment was done with 10 µM Sorafenib tosylate (Sellekchem) for up to 2 h.

### Nanobody immobilization on NHS-Sepharose matrix

2 mL of purified Miro1-Nbs at 1 mg/mL concentration in phosphate-buffered saline (PBS) were immobilized on 1 mL NHS-Sepharose (GE-Healthcare) according to the manufacturer’s protocol.

### Sortase labelling of Nanobodies

Sortase A pentamutant (eSrtA) in pET29 expression vector, a gift from David Liu (Addgene plasmid # 75144) was expressed and purified as described [78]. The substrate peptide H-Gly-Gly-Gly-propyl-azide (sortase substrate) was custom synthesized by Intavis AG. For sortase labelling, 50 μM Nb, 250 μM sortase substrate peptide dissolved in sortase buffer (50 mM Tris, pH 7.5, and 150 mM NaCl) and 10 μM sortase were mixed in coupling buffer (sortase buffer with 10 mM CaCl_2_) and incubated for 4 h at 4 °C. Uncoupled Nb and sortase were depleted by IMAC. Unbound excess of unreacted peptide was removed using Zeba Spin Desalting Columns (ThermoFisher Scientific). Azide-coupled Nbs were then labelled by SPAAC (strain-promoted azide-alkyne cycloaddition) click chemistry reaction with 2-fold molar excess of Alexa-Fluor conjugated dibenzocyclooctyne (DBCO-AF647, Jena Bioscience) for 2 h at 25 °C. For generation of the bivalent nanotraps, 1 mg of azide coupled bivalent Nb was incubated with 0.5 mL DBCO-Agarose (Jena Bioscience) slurry for 4 h at 25 °C. The beads were centrifuged at 2700 x g for 2 min and the supernatant was removed. Samples of the input and flow-through fractions were analysed by SDS-PAGE according to standard protocol. The Nb-coupled beads were washed thrice with 2.5 mL PBS and stored in 1 mL PBS.

### Immunoprecipitation

2 x 10^6^ HEK 293 cells were seeded in 100 mm culture dishes (Corning) and cultivated for 24 h. For the pulldown of endogenous Miro1, the cells were harvested and lysed after 24 h. For the pulldown of GFP-Miro1 or GFP, the cells were subjected to plasmid DNA transfection with equal amounts of expression vectors and cultivated for 24 h. Subsequently, cells were washed in PBS (pH 7.4) and harvested, snap-frozen in liquid nitrogen, stored at -20 °C or thawed for immediate use. Cell pellets were homogenized in 200 µL lysis buffer (50 mM Tris/HCl pH 7.5, 150 mM NaCl, 1 mM EDTA, 0.5% Triton X-100, 1 mM PMSF, 1 µg/mL DNaseI, 2.5 mM MgCl_2_, 1 x protease inhibitor cocktail (Serva)) by passing through needles of decreasing gauge and intermittent vortexing for 60 min on ice. Lysates were clarified by subsequent centrifugation at 18,000 x g for 15 min at 4 °C. The supernatant was adjusted with dilution buffer (50 mM Tris/HCl, 150 mM NaCl, 0.5 mM EDTA, 1 mM PMSF) to 500 µL. 5 μL (1%) was added to 2 x SDS-containing sample buffer (60 mM Tris/HCl, 2% (w/v) SDS, 5% (v/v) 2-mercaptoethanol, 10% (v/v) glycerol, 0.02% bromphenol blue; referred to as input). For immunoprecipitation, 40 – 80 μl of sepharose-coupled Miro1 Nbs (nanotraps) were added to the protein solution and incubated for 16 h on an end over end rotor at 4 °C. As a positive control, 2.5 µg of rabbit anti- Miro1 antibody (# PA5-42646, ThermoFisher) was added to the protein solution and incubated under the same conditions. For pulldown of immunocomplexes, 40 μl of an equilibrated mixture of protein A/G-Sepharose (Amersham Biosciences) were added, and incubation continued for 4 h. As a negative control, a non-related Nb (PepNb) immobilized on sepharose beads was used. After centrifugation (2 min, 2700 x g, 4 °C) supernatant was removed and the bead pellet was washed three times in 0.5 mL dilution buffer. On the third wash, the beads were transferred to a pre-cooled 1.5 mL tube (Eppendorf), resuspended in 2 x SDS-containing sample buffer and boiled for 10 min at 95 °C. Samples (1% input, 20% bound) were analysed by SDS-PAGE followed by Western blotting according to standard procedures. Immunoblots were probed with the following primary antibodies: anti-Miro1 (clone 4H4, Sigma-Aldrich), anti-GFP (clone 3H9, ChromoTek) and anti-GAPDH (Santa Cruz) antibody as a negative control to detect unspecific binding to the nanotraps. Full scans of western blots from all immunoprecipitation experiments are included in **Supplementary Figure 12**.

### Immunofluorescence

HeLa cells were seeded at 1 x 10^4^ per well in μClear 96 well plates (Greiner). Next day, the cells were transfected with plasmids coding for GFP-Miro1 or a GFP-tagged non-related protein. 24 h post transfection, the cells were washed twice with PBS and fixed with 4% (w/v) paraformaldehyde (PFA) in PBS for 10 min at RT and blocked with 5% BSA in TBST for 30 min at RT. Incubation with purified Miro1 Nbs or AF647 conjugated Nbs (100 – 200 nM in 5% BSA in TBST) or rabbit anti-Miro1 antibody (# NBP1-89011, Novus Biologicals) was performed overnight at 4 °C. Unbound nanobodies were removed by three washing steps with TBST. Unlabelled Miro1 Nbs were detected by addition of a Cy5-conjugated Goat Anti-Alpaca IgG (Jackson Immunoresearch) according to manufacturer’s guidelines. For Miro1 antibody detection, an AF647 conjugated goat anti-rabbit secondary antibody (Invitrogen) was used. Nuclei were subsequently stained with 4’, 6-diamidino-2-phenylindole (DAPI, Sigma-Aldrich) and images were acquired immediately afterwards with an ImageXpress™ Micro Confocal High Content Screening system (Molecular Devices) at 40x magnification.

### Liquid chromatography-MS analysis

Miro1 pull-down samples, by application of the mono- and bivalent M41 and M114 nanotraps, were compared to control nanotraps in three technical replicates. Proteins were purified by SDS- PAGE (4-12% NuPAGE tris gel (Invitrogen) for 7 min at 200 V. Coomassie stained protein gel pieces were excised and applied to tryptic digestion as described previously [79]. Samples were measured on an Exploris480 mass spectrometer (Thermo Fisher Scientific) online-coupled to an Easy-nLC 1200 UHPLC (Thermo Fisher Scientific). Peptides were separated on an in-house packed (ReproSil-Pur C18-AQ 1.9 μm silica beads (Dr Maisch GmbH, Ammerbuch, Germany)), 20 cm analytical HPLC column (75 μm ID PicoTip fused silica emitter (New Objective, Berks, UK)). Peptides were eluted with a 36 min gradient, generated by solvent A (0.1% formic acid) and solvent B (80% Acetonitrile in 0.1% formic acid) at a flow rate of 200 nL/min at 40°C. Nanospray ionization at 2.3 kV together with a capillary temperature of 275°C was applied for peptide ionization. Full MS spectra were acquired at resolution 60k within a scan range of 300-1750 m/z and tandem MS (MS/MS) spectra were acquired at 15k resolution. Maximum Injection Time Mode and automated control target were set to Auto and Standard respectively for full MS and MS/MS scans. The 20 most intense peptides with multiple charge were selected for MS/MS sequencing by higher-energy collisional dissociation (HCD) with a dynamic exclusion of 30 s.

### Mass Spectrometry Data analysis

Raw data files were processed using the MaxQuant software suit (version 2.0.3.0)[80]. Spectra were searched against Uniprot *Homo sapiens* database (released 11.12.2019, 96,817 entries), *Vicugna pacos* specific nanotraps and commonly observed contaminants. Peptide mass tolerance was set to 4.5 ppm for MS and to 20 ppm for MS/MS. Peptide and protein false discovery rate was set to 1%. Methionine oxidation and protein N-terminus acetylation were selected as variable modification, while carbamidomethylation on Cysteine was defined as fixed modification. A maximum of two missed cleavages were accepted for specific trypsin digestion mode. For label-free quantification a minimum number of two ratio count was requested. Intensity based absolute quantification was enabled. Statistical analysis was performed with the Perseus software suit (version 1.6.15.0). First, contaminants, reversed and proteins only identified by site proteins were filtered out. Only proteins present in two out of three replicates of each nanotrap were allowed for downstream significance testing. Significantly enriched proteins were determined by t-test as Class A with S0 set to 0.1 and FDR threshold ≤ 0.01 or Class B with S0 set to 0.1 and FDR threshold ≤ 0.05. Additional graphical visualization were performed in the R environment (version 4.1.1). For Miro1 interactome analysis the top 50 protein interactors were classified based on the direct or indirect interaction with Miro1 and the confidence score derived from STRING database. As class 1 interactors, proteins were assigned that are direct interactors with Miro1, and had a confidence score greater than 0.9. Proteins with a direct interaction with Miro1 and a confidence score smaller than 0.9 were annotated as class 2 interactors. Class 3 interactors were annotated based on indirect Miro1 interaction and a confidence score greater than 0.9.

### Microscopy and time lapse imaging

8 x 10^3^ U2OS or 1 x 10^4^ HeLa cells/well were plated in a black µclear 96-well plate (Greiner). 24h after plating, cells were transiently co-transfected with plasmids coding for GFP-Miro1 and M41-Cb, M85-Cb or M114-Cb. The next day, the medium was replaced by live-cell visualization medium DMEM^gfp-2^ (Evrogen) supplemented with 10% FBS and 2 mM L-glutamine. For the time-lapse acquisition, U2OS cells transiently expressing GFP-Miro1 and M114-CB or mito-mKate2 were either treated with 10 µM Sorafenib tosylate or DMSO in live-cell visualization medium and imaged every 15 min for up to 2 h. Images were acquired under standard conditions with the ImageXpress™ Micro Confocal High Content Screening system (Molecular Devices) at 40x magnification.

### Image segmentation and analysis

For the targeted Miro1 degradation experiments, 8 x 10^3^ - 1 x 10^4^ wildtype HeLa cells or HeLa cells transiently expressing Fbox-Nb-IRES-tRFP-NLS constructs were fixed and permeabilized in a black µclear 96-well plate (Greiner). Immunofluorescence staining of Miro1 was performed as previously described and images were acquired immediately with an ImageXpress™ micro XL system (Molecular Devices) at 40x magnification. Image analysis was performed with MetaXpress software (64 bit, 6.2.3.733, Molecular Devices). Fluorescence images comprising a statistically relevant number of cells (>200 cells) were acquired for each construct. For quantitative fluorescence analysis, the mean Miro1 fluorescence in the cytosol was determined. Using the Custom Module Editor (version 2.5.13.3) of the MetaXpress software, we established an image segmentation algorithm that identifies areas of interest based the parameters of size, shape, and fluorescence intensity above local background. For nuclear segmentation, DAPI stained nuclei or nuclear localized TagRFP in transfected cells were defined as selection criteria. To segment the cytosolic compartment of the cell, Miro1 fluorescence signals were used to generate a segmentation mask. The average Miro1 fluorescence intensities in whole cells were determined for each image followed by subtraction of background fluorescence. The resulting values from the transfected cells were normalized to the non-transfected control. Standard errors were calculated for three independent replicates and student’s *t* test was used for statistical analysis.

### Analyses and Statistics

Graph preparation and statistical analysis was performed using the GraphPad Prism Software (Version 9.0.0 or higher).

## Author contributions

F.O.F. and U.R. conceived the study and analyzed the data. F.O.F., P.D.K., T.R.W., B.T., T.F., and G.J. perform all cellular and biochemical experiments including imaging studies. N.B. provided recombinant Miro1. S.N. and A.S. immunized the alpaca. K.Z. and B.M. performed and analyzed mass spectrometry experiments. F.O.F. and U.R. wrote the manuscript with the help of all authors.

## Acknowledgements

This research was supported by the German Research Foundation (DFG) through RTG 2364 “MOMbrane” to F.O.F., K.Z., B.M and U.R. The authors also gratefully acknowledge the Ministry of Science, Research and Arts of Baden-Württemberg (V.1.4.-H3-1403-74) for financial support.

## Supplementary information

### Supplementary Data

**Supplementary Figure 1.**
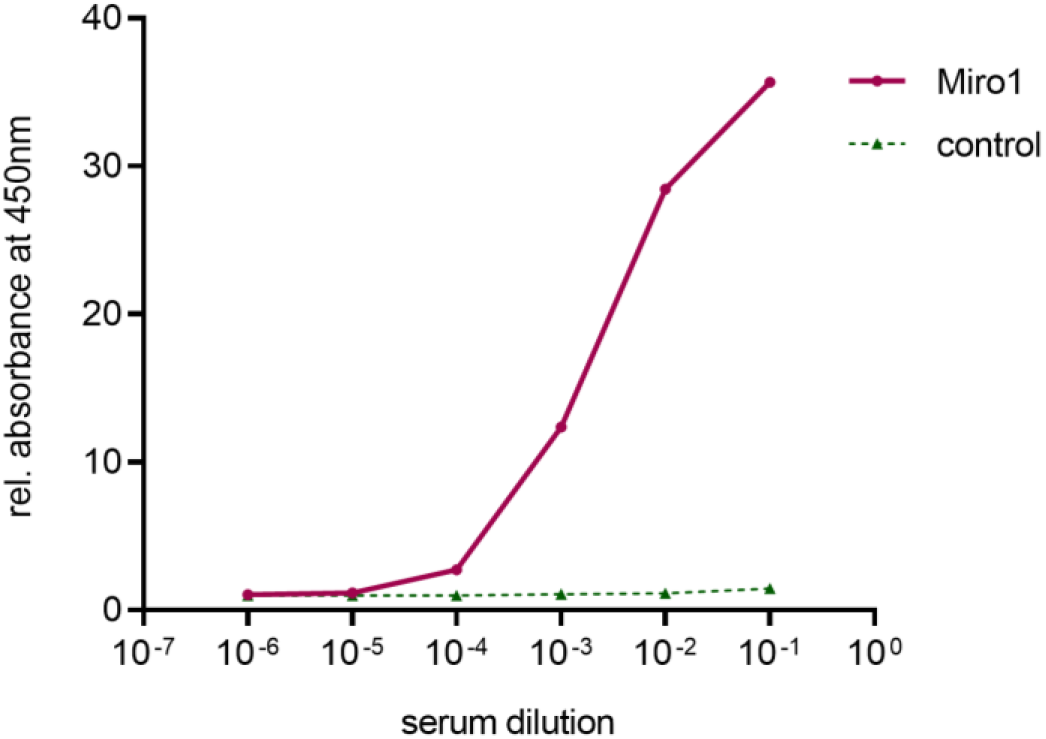
Analysis of seroconversion upon vaccination with Miro1. To monitor an immune response upon vaccination, a serum sample was from the immunized Alpaca (Vicugna pacos) on day 63 after starting immunization. Formation of Miro1 specific antibodies by the animal was measured in a serum ELISA at indicated dilutions in multiwall plates either coated with hMiro1 or bovine serum albumin (BSA) as negative control. Bound antibodies were detected using an anti-heavy chain antibody secondary antibody labelled with horse radish peroxidase. Obtained ELISA signals for hMiro1 were normalized to signals obtained for the negative control.

**Supplementary Figure 2.**
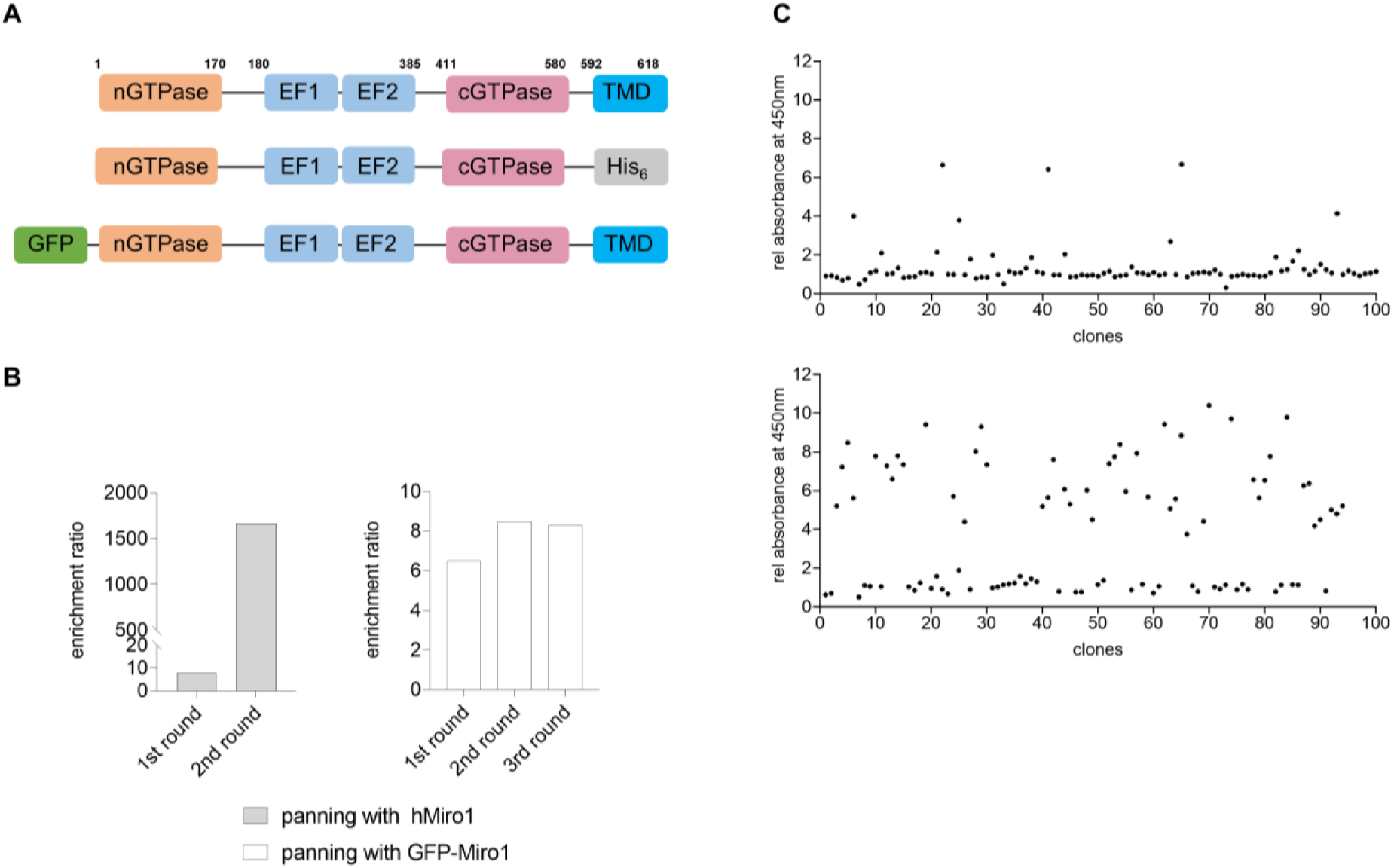
Enrichment and selection of Miro1 nanobodies (Nbs) by phage display and phage ELISA. (A) Illustration of wildtype Miro1 and recombinant Miro1 constructs used for phage display. (B) Bar charts showing the enrichment of Miro1-Nb phages after two iterative panning rounds against bacterial expressed hMiro1 (left panel, grey bars) and three iterative panning rounds against GFP-Miro1 (right panel, white bars). (C) Phage ELISA profile of 100 eluted phage clones tested for binding to hMiro1 (top panel) or GFP-Miro1 (lower panel). Signal intensities were normalized to a signal obtained for BSA used as negative control.

**Supplementary Figure 3.**
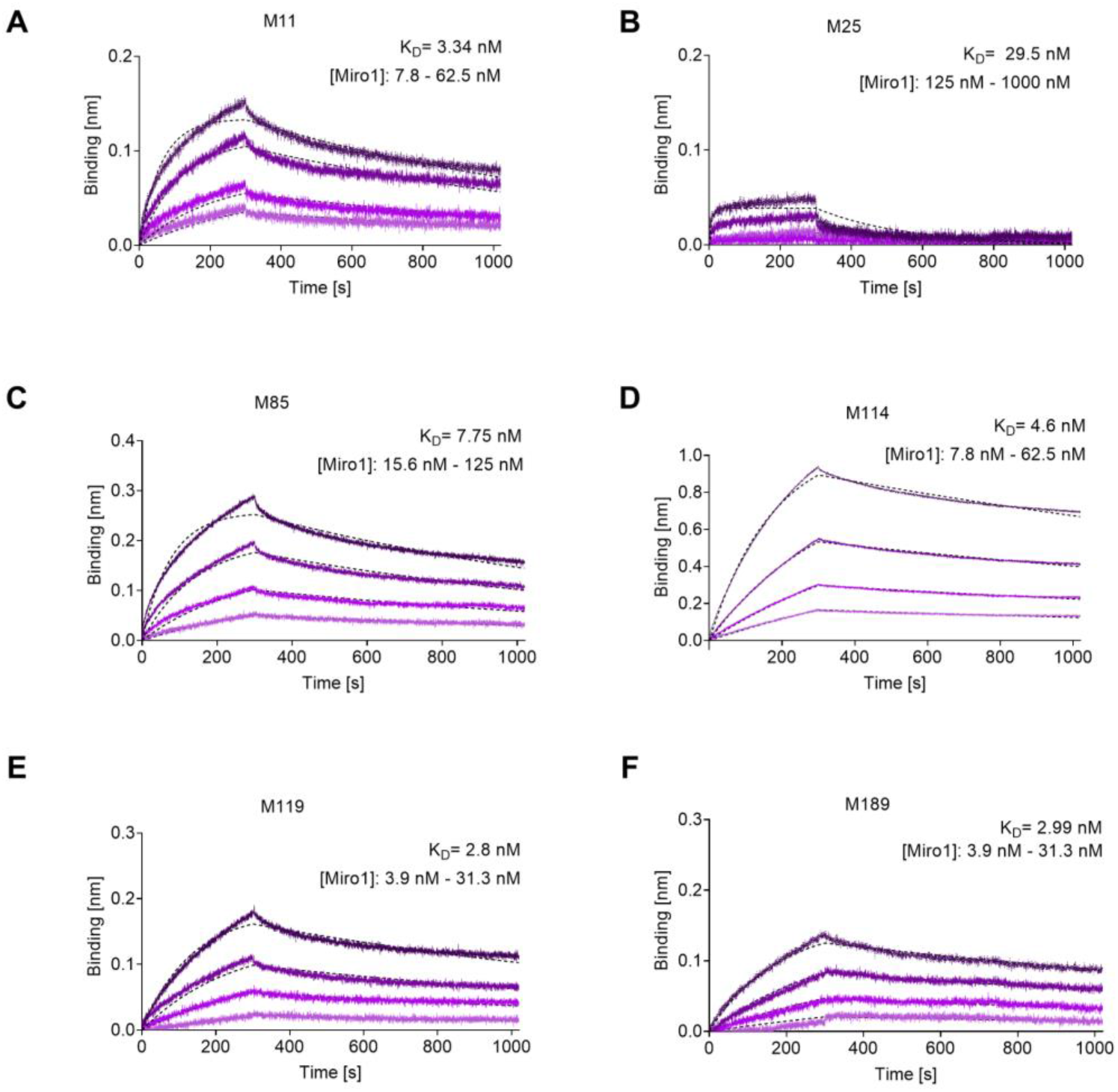
Affinities of identified Miro1-Nbs. Affinities of Miro1-Nbs were analysed by biolayer interferometry based affinity measurements. Miro1-Nbs were biotinylated and immobilized on streptavidin sensors. Kinetic measurements were performed by using four concentrations of hMiro1. The sensograms of hMiro1 on M11 (**A**), M25 (**B**), M85 (**C**), M114 (**D**), M119 (**E**) and M189 (**F**) at indicated concentrations (illustrated with increasingly darker shades from low to high concentration) are shown and global 1:1 fits are illustrated as dashed lines. A summary of the affinities (K_D_), association constants (K_ON_) and dissociation constants (K_OFF_) determined for all seven Miro1-Nbs are shown in Figure 1C**, Table 1**.

**Supplementary Figure 4.**
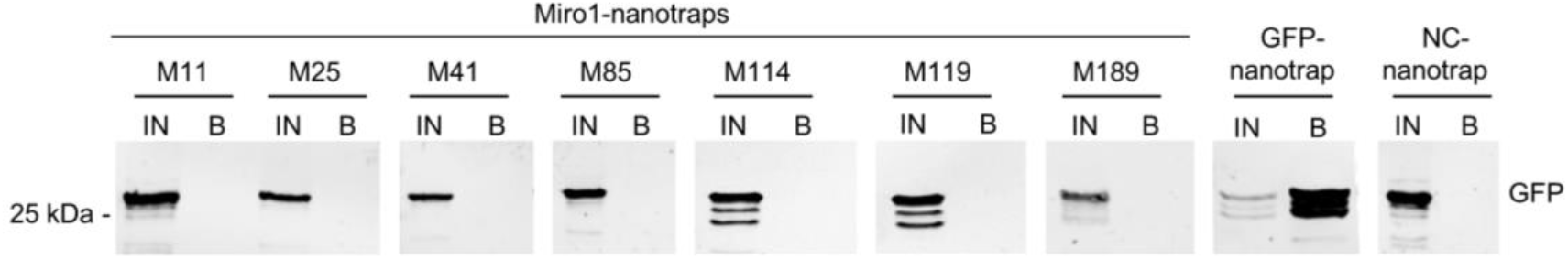
Immobilized Miro1-Nbs (nanotraps) do not bind GFP. Soluble protein fraction of HEK293 cells transiently expressing GFP as control antigen, was adjusted to 2 mg/mL and incubated with equal amounts of nanotraps. Input (IN, 1% of total) and bound (20% of total) fractions were subjected to SDS-PAGE followed by immunoblot analysis using antibodies specific for GFP. As positive control GFP-nanotrap and as negative a non-specific (NC) nanotrap were used.

**Supplementary Figure 5.**
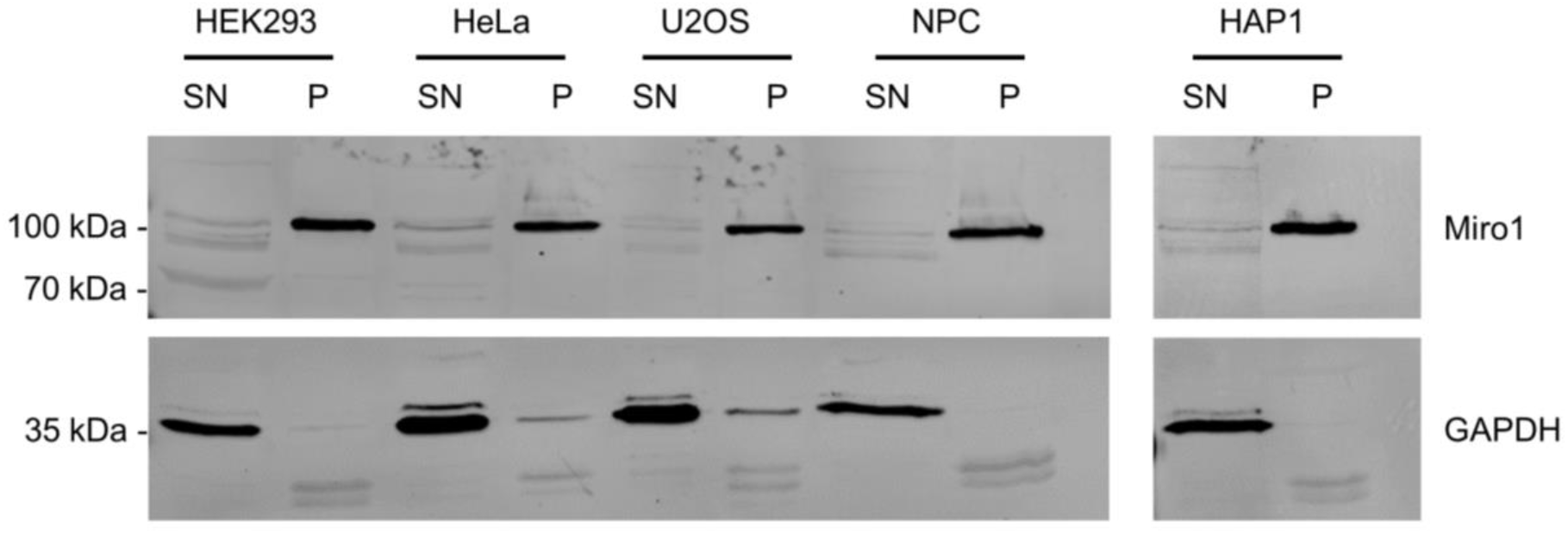
Endogenous Miro1 retains in the insoluble protein fraction in different cell lines. Representative western blot analysis of 20 µg of soluble (SN) and insoluble/pellet (P) fractions of HEK293, HeLa, U2OS, HAP1 and neuroprogenitor (NPC) cells. Shown is a representative example of an immunoblot after cell lysis using 0.5% NP-40 in the lysis buffer. The upper part of the blots were detected with anti-Miro1 antibody. Detection of GAPDH with an anti-GAPDH antibody was used as lysis and loading control.

**Supplementary Figure 6.**
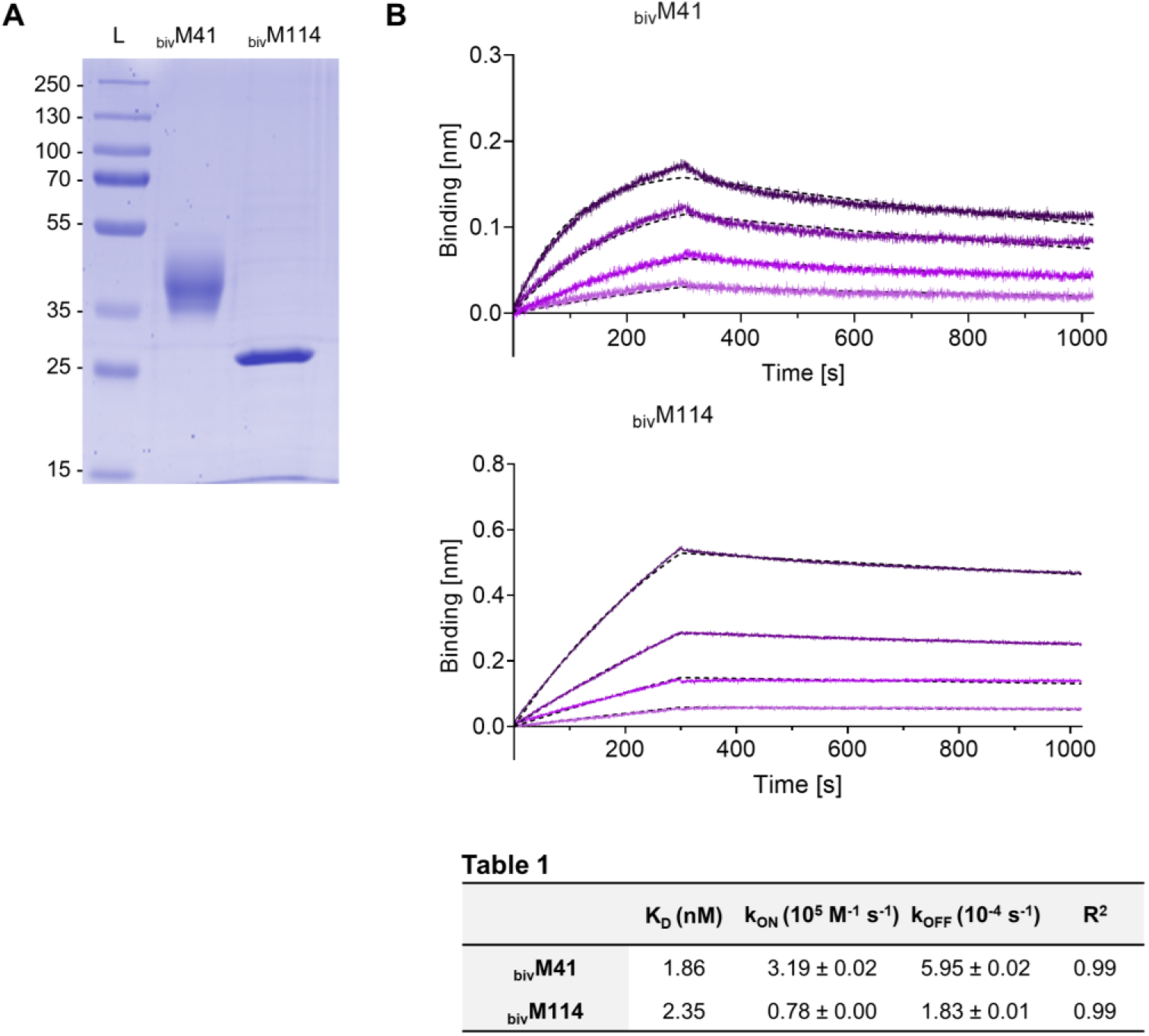
Recombinant expression, purification and characterization of bivalent M41- and M114-Nbs. (A) Coomassie stained SDS-PAGE of 2 µg _biv_M41- and _biv_M114-Nbs purified from ExpiCHO™ cells is shown. (B) Affinities of bivalent Miro1-Nbs were analysed by biolayer interferometry (BLI) based affinity measurements. Bivalent Miro1-Nbs were biotinylated and immobilized on streptavidin sensors. Kinetic measurements were performed by using four concentrations of hMiro1 ranging from 3.9 nM – 31.3 nM (illustrated with increasing concentrations in darker shades). The sensograms of purified Miro1 on _biv_M41-Nb (top) and _biv_M114-Nb (bottom) are shown and global 1:1 fits are illustrated as dashed lines. The table summarizes affinities (K_D_), association (k_ON_) and dissociation constants (k_OFF_), and coefficient of determination (R^2^) determined for both bivalent Nbs.

**Supplementary Figure 7.**
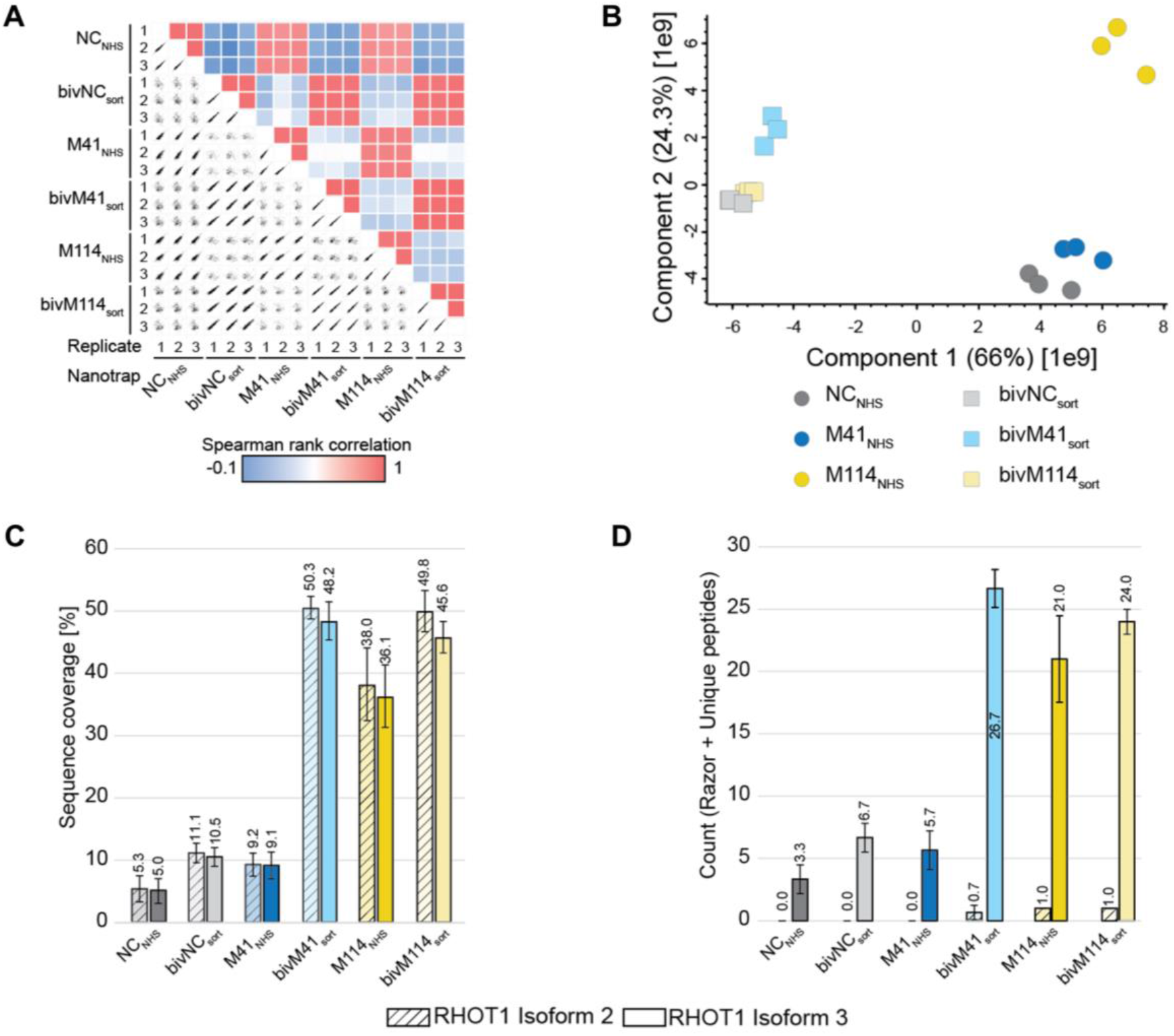
Enrichment efficiency of bivalent nanotraps. (A) Multi correlation between replicates and nanotraps. High Spearman rank correlation between replicates and mono- or bivalent nanotraps. (B) Principle component analysis (PCA) reflects highest similarity between replicates. 66% of variance between samples explained by nanotrap valency. (C) Identification of Miro1 peptides (razor and unique) for Miro1 isoforms. Averaged across three replicates. (D) Averaged sequence coverage of Miro1 higher for bivalent nanotraps

**Supplementary Figure 8.**
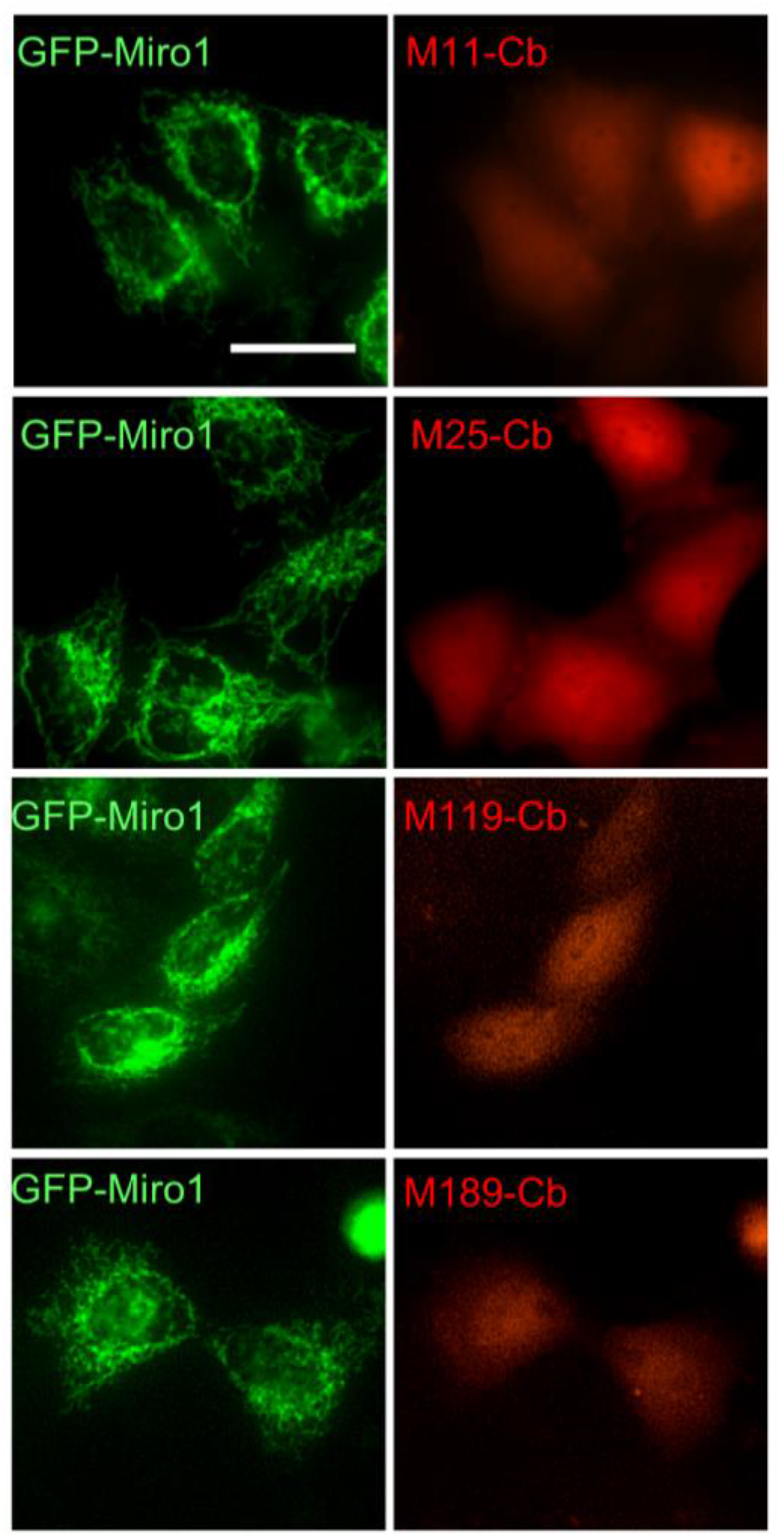
Detection of intracellular binding capacities of Miro1-Cbs. Representative fluorescence images of live HeLa cells transiently expressing GFP-Miro1 (left column) in combination with TagRFP-labelled Miro1-Cbs (right column). Scale bar 20 µm.

**Supplementary Figure 9.**
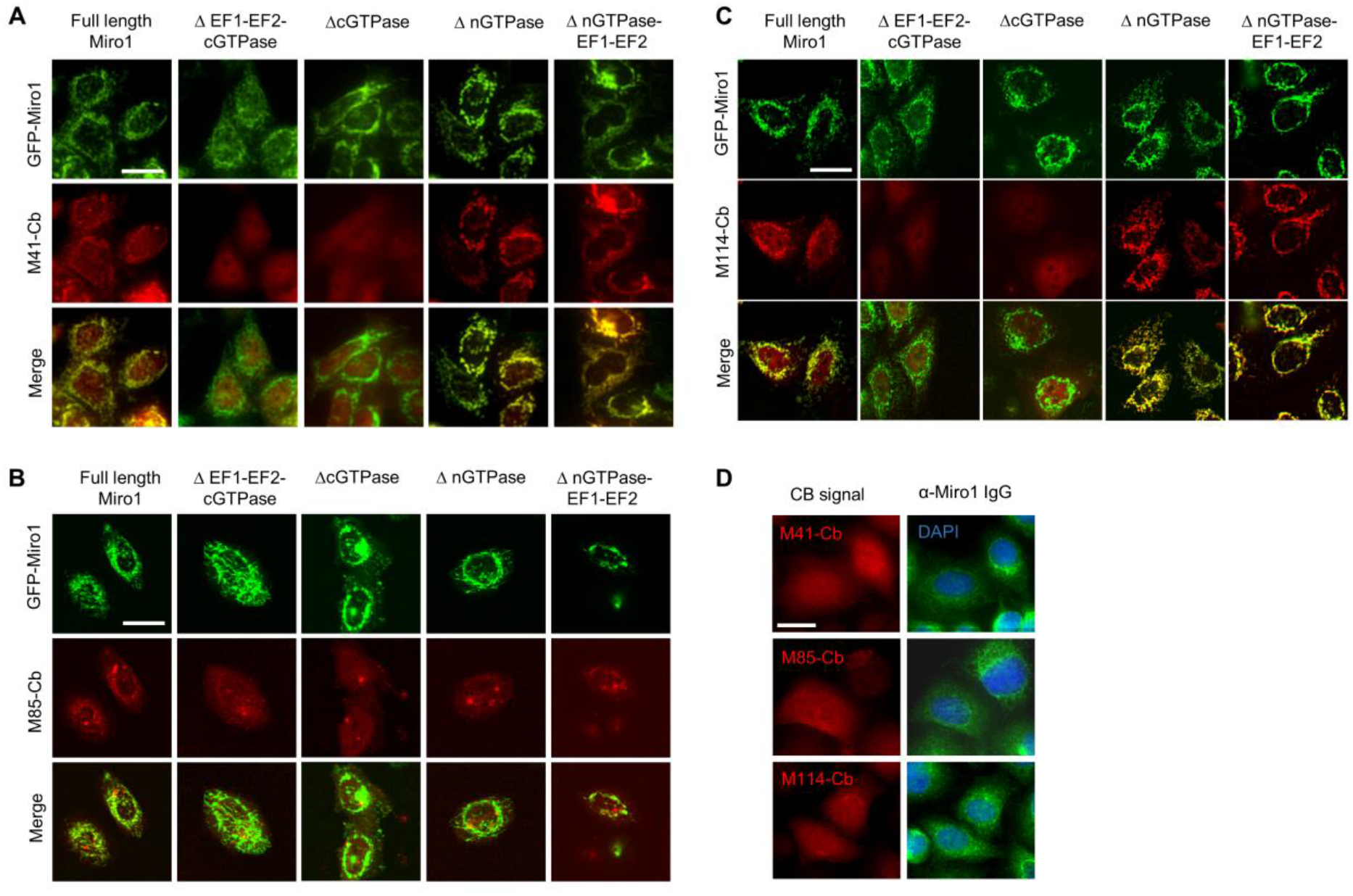
Intracellular characterization of domain specific binding of selected Miro1-Cbs. Representative fluorescent images of live HeLa cells transiently expressing GFP-Miro1 and indicated GFP-tagged Miro1 domain deletion constructs (top row) in combination with TagRFP-labelled M41-Cb (**A**), M85-Cb (**B**) or M114-Cb (**C**) (middle row). Scale bar 20 µm. (**D**) Immunofluorescence detection of endogenous Miro1 in HeLa cells expressing M41-, M85- and M114-Cb (left panel) using an anti-Miro1 antibody (right panel). Scale bar 25 µm.

**Supplementary Figure 10.**
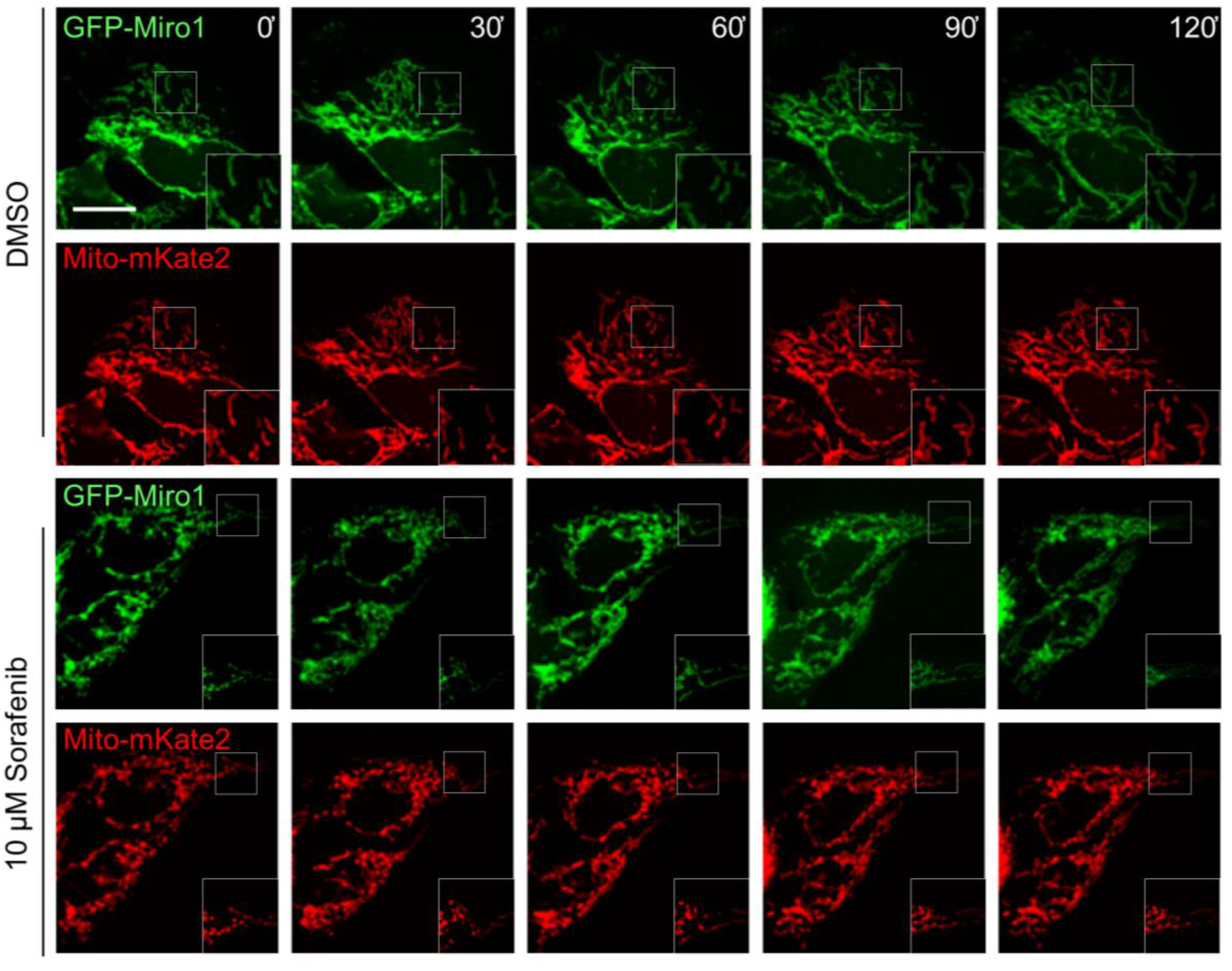
Live-cell imaging of mitochondrial phenotypes. Time-lapse microscopy of U2OS cells transiently expressing GFP-Miro1 and mitoMkate2 (as a mitochondrial marker). To visually track morphological mitochondrial changes, cells were treated with either DMSO as a control (top two rows) or 10 µM Sorafenib (bottom two rows) followed by time-lapse imaging over a 2 hour period. Shown are representative images of three biological replicates. Scale bar 25 µm. Squares at the bottom right represent enlargements of the selected image section.

**Supplementary Figure 11.**
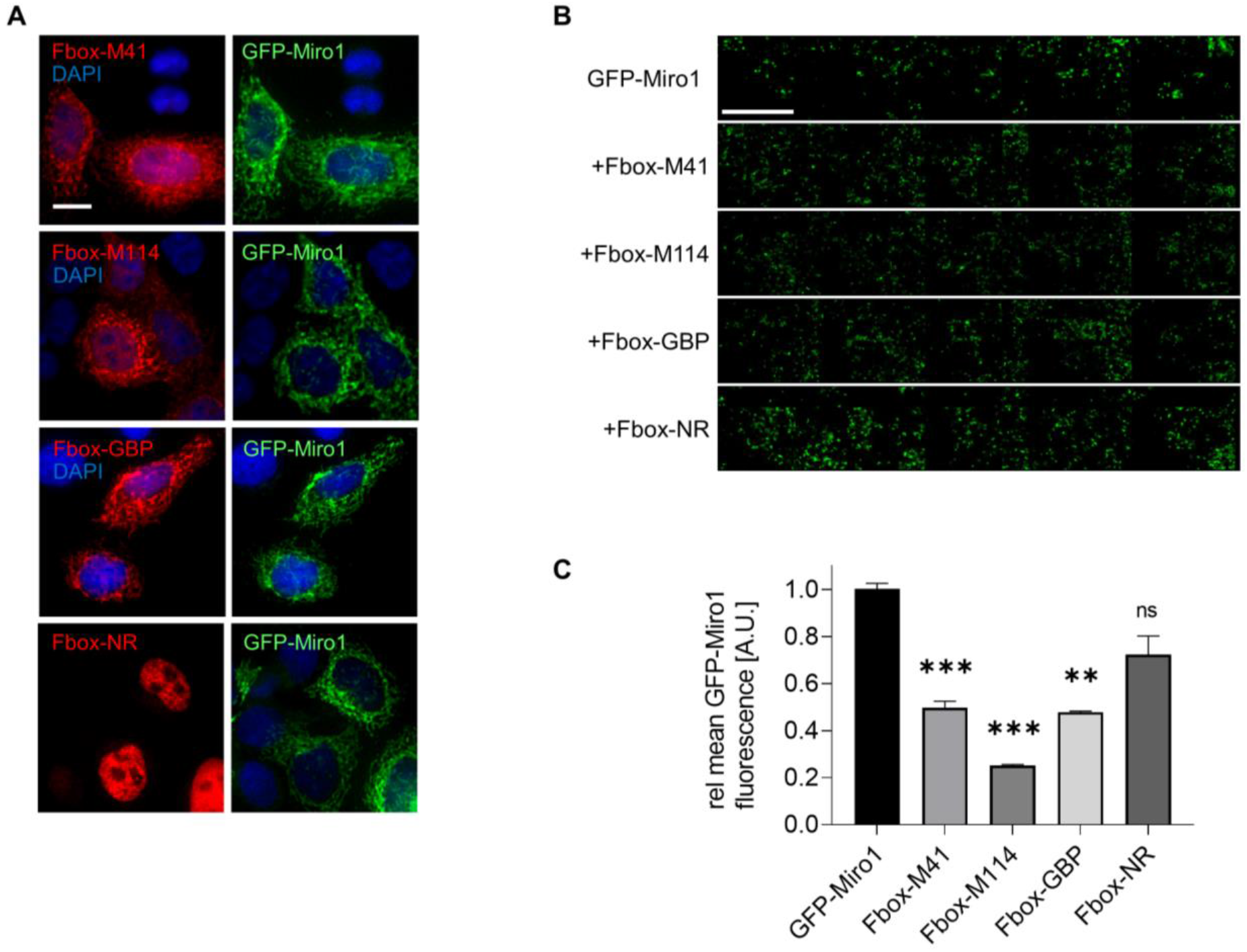
Targeted degradation of GFP-Miro1 by Fbox-Nb-based degrons in live cells. (A) Representative confocal images of HeLa cells transiently co-expressing GFP-Miro1 and indicated Miro1-specific Fbox-Nbs (Fbox-M41, Fbox-M114) or a non-related Fbox-Nb (Fbox- NR) construct are shown. For quantitative IF analysis, cells were fixed and permeabilized 24 h after transfection followed by staining with Cy5 conjugated anti-VHH antibody and DAPI. Fbox- Nb expressing cells were subjected to automated image analysis and quantification as described in Material and Methods. Scale bar 20 µm. (B) Representative fluorescence thumbnail images of HeLa cells coexpressing GFP-Miro1 and Fbox-Nb constructs. Scale bar 1 mm. (C) Mean GFP-Miro1 fluorescence intensity from HeLa cells co-expressing GFP-Miro1 and Fbox-Nbs constructs were determined by quantitative fluorescence imaging. Fluorescence intensity values were calculated from three samples (n= 3; >500 cells) and normalized to the GFP-Miro1 signal intensity (set to 1). Positive control; GFP-specific Fbox-Nb (Fbox-GBP), negative control; non-related Fbox-Nb construct (Fbox-NR). Data are represented as mean ± SEM. For statistical analysis, student‘s t-test was performed, **p < 0.01, ***p < 0.001.

**Supplementary Figure 12.**
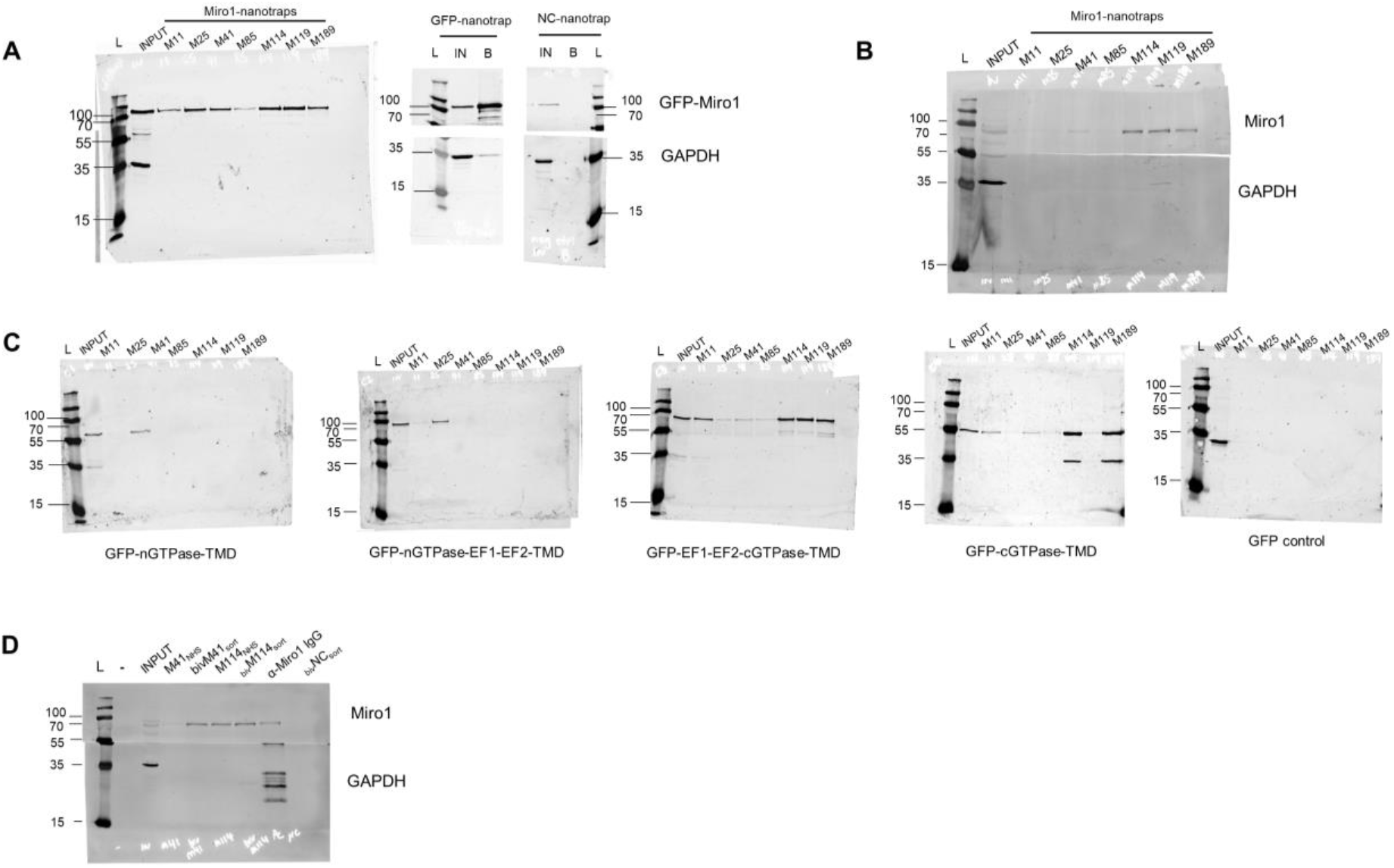
Western blot data. (A) Full size of Western blots shown in Figure 2A, top halves stained with anti-GFP antibody, bottom halves stained with anti-GAPDH antibody (B) Full size of Western blots shown in Figure 2B, top half detected by anti-Miro1 antibody, bottom half by anti-GAPDH antibody. (C) Full size of Western blots shown in Figure 3B, detection with anti-GFP antibody. (D) Full size of Western blots shown in Figure 5, **t**op half detected by anti-Miro1 antibody and the bottom half by anti-GAPDH antibody.

**Supplementary Table 1.**
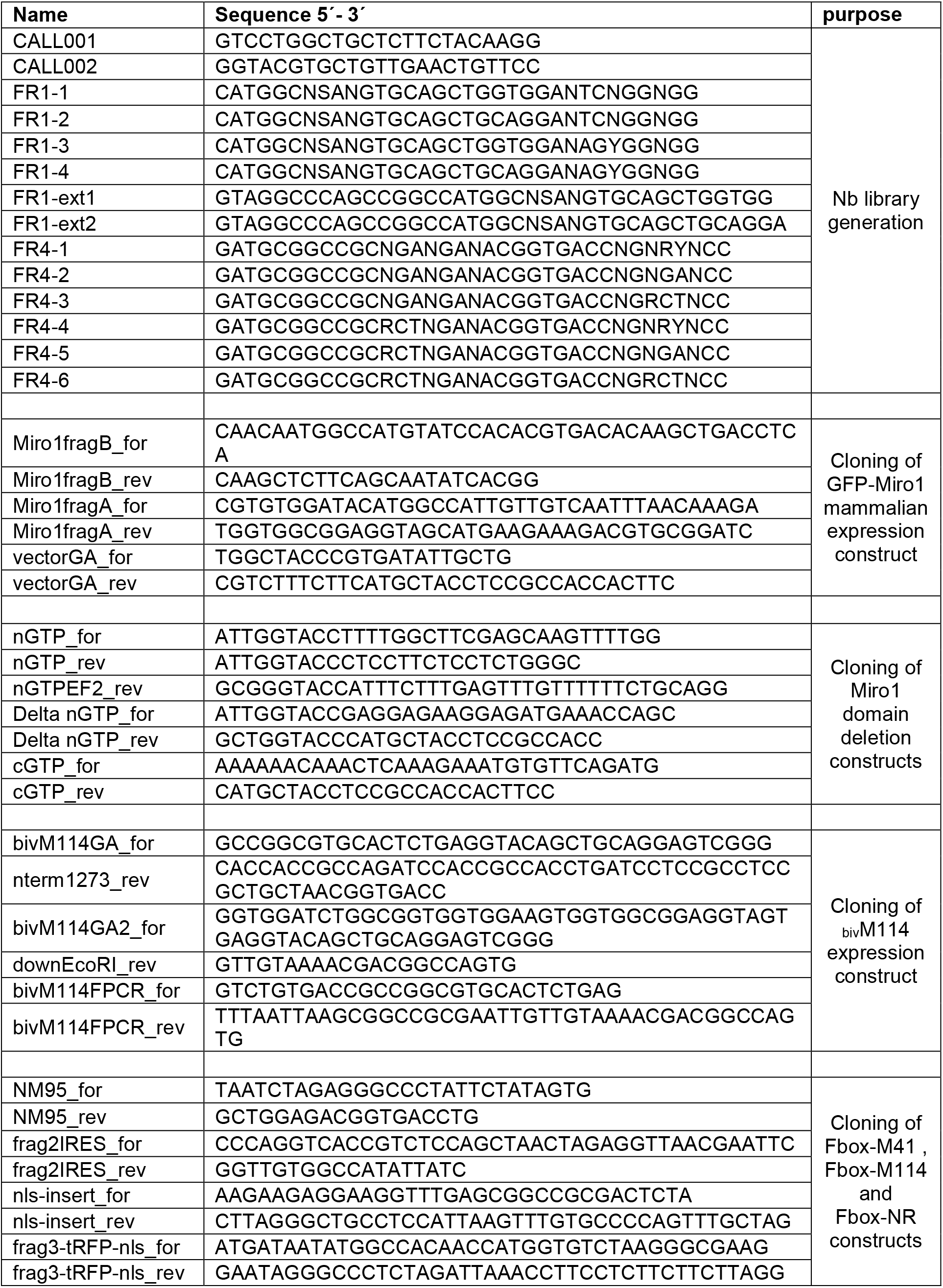
List of oligonucleotides used in this study.

**Supplementary Table 2.**
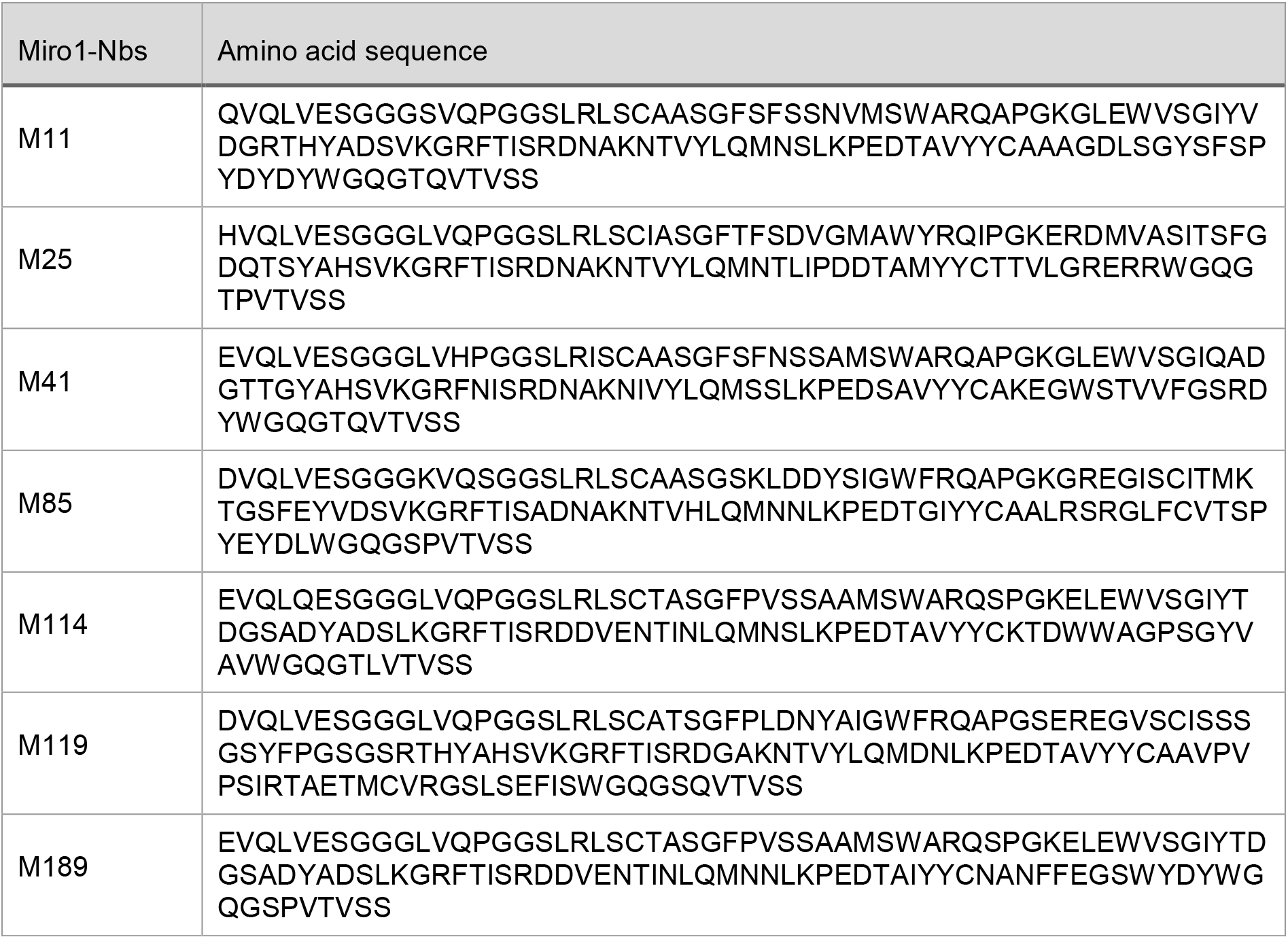
Amino acid sequences of all selected Miro1-Nbs.

